# Effect of selective lesions of nucleus accumbens μ-opioid receptor-expressing cells on heroin self-administration in male and female rats: a study with novel *Oprm1-Cre* knock-in rats

**DOI:** 10.1101/2022.11.02.514895

**Authors:** Jennifer M. Bossert, Carlos A. Mejias-Aponte, Thomas Saunders, Lindsay Altidor, Michael Emery, Ida Fredriksson, Ashley Batista, Sarah M. Claypool, Kiera E. Caldwell, David J. Reiner, Jonathan J. Chow, Matthew Foltz, Vivek Kumar, Audrey Seasholtz, Elizabeth Hughes, Wanda Filipiak, Brandon K. Harvey, Christopher T. Richie, Francois Vautier, Juan L. Gomez, Michael Michaelides, Brigitte L. Kieffer, Stanley J. Watson, Huda Akil, Yavin Shaham

**Author notes:** **Correspondence:** Jennifer M. Bossert or Yavin Shaham. <u>Author Contributions:</u> The authors made significant contributions to the paper by either being involved in the creation or validation of the new rat line.

## Abstract

The brain µ-opioid receptor (MOR) is critical for the analgesic, rewarding, and addictive effects of opioid drugs. However, in rat models of opioid-related behaviors, the circuit mechanisms of MOR-expressing cells are less known because of a lack of genetic tools to selectively manipulate them. We introduce a CRISPR-based *Oprm1*-Cre knock-in transgenic rat that provides cell-type specific genetic access to MOR-expressing cells. After performing anatomical and behavioral validation experiments, we used the *Oprm1*-Cre knock-in rats to study the role of nucleus accumbens (NAc) MOR-expressing cells in heroin self-administration in male and female rats.

Using RNAscope, autoradiography, and fluorescence *in situ* hybridization chain reaction (HCR-FISH), we found no differences in *Oprm1* expression in NAc, dorsal striatum (DS), and dorsal hippocampus, or MOR receptor density (except DS) or function between *Oprm1*-Cre knock-in rats and wildtype littermates. HCR-FISH assay showed that *iCre* is highly co-expressed with *Oprm1* (95-98%). There were no genotype differences in pain responses, morphine analgesia and tolerance, heroin self-administration, and relapse-related behaviors. We used the Cre-dependent vector AAV1-EF1a-Flex-taCasp3-TEVP to lesion NAc MOR-expressing cells and report sex-specific effects: the lesions decreased acquisition of heroin self-administration in male *Oprm1*-Cre rats and had a stronger inhibitory effect on the effort to self-administer heroin in female *Oprm1*-Cre rats.

The validation of an *Oprm1*-Cre knock-in rat enables new strategies for understanding the role of MOR-expressing cells in rat models of opioid addiction, pain-related behaviors, and other opioid-mediated functions. Our initial mechanistic study with these rats suggests a sex-specific role of NAc MOR-expressing cells in heroin self-administration.

**Significance statement:** The brain µ-opioid receptor (MOR) is critical for the analgesic, rewarding, and addictive effects of opioid drugs. However, in rat models of opioid-related behaviors, the circuit mechanisms of MOR-expressing cells are less known because of a lack of genetic tools to selectively manipulate them. We introduce a CRISPR-based *Oprm1*-Cre knock-in transgenic rat that provides cell-type specific genetic access to brain MOR-expressing cells. After performing anatomical and behavioral validation experiments, we used the *Oprm1*-Cre knock-in rats to show a potential sex-specific role of nucleus accumbens MOR-expressing cells in heroin self-administration. The new *Oprm1*-Cre rats can be used to study both the general and sex-specific role of brain MOR-expressing cells in animal models of opioid addiction, pain-related behaviors, and other opioid-mediated functions.

## Introduction

The µ-opioid receptor (MOR) is expressed in many brain areas (Akil et al., 1984; Mansour et al., 1995a; Emery and Akil, 2020). Activation of MORs mediates diverse effects of opioid agonists like heroin, morphine, and fentanyl, including analgesia, tolerance, and self-administration in mice, rats, monkeys, and humans, as well as addiction liability of opioid drugs in humans (Jaffe, 1990; Darcq and Kieffer, 2018).

During the last decade, investigators have developed transgenic mouse models that allow for the investigation of circuit mechanisms of MOR-expressing cells in different brain regions in the behavioral and physiological effects of opioid drugs. These include a knock-in MOR-mCherry mouse line to map MOR protein expression throughout the brain (Gardon et al., 2014) and a mouse line with a floxed *Oprm1* gene (the gene encoding MOR) that allows for selective deletion of the receptor (Weibel et al., 2013; Charbogne et al., 2017). More recently, Bailly et al. (2020) introduced an *Oprm1*-Cre knock-in mouse line that allows for *in vivo* manipulation of activity of MOR-expressing cells in the brain to study their causal role in the behavioral and physiological effects of opioid drugs. In this *Oprm1*-Cre mouse line, a cDNA encoding a T2A cleavable peptide and Cre recombinase was fused to enhanced green fluorescent protein (eGFP), and the genetic construct was inserted in frame downstream of the *Oprm1* coding sequence.

To date, the Cre line technology has not been applied to study the role of MOR-expressing cells and projections in opioid analgesia and self-administration in the rat. Mice provide a good model organism to study circuit mechanisms of unconditioned and simple conditioned behaviors related to opioid analgesia and reinforcement. However, it has been difficult to reliably study circuit mechanisms of opioid self-administration and relapse-related behaviors in this species due to technical limitations (small veins and difficulties in maintaining catheter patency) and limited repertoire of sophisticated learned behaviors. For example, established rat behavioral phenomena like incubation of drug craving (time-dependent increase in drug seeking during abstinence) and drug priming-induced reinstatement after extinction (Shaham et al., 2003; Wolf, 2016) are not readily observed in mouse models (Highfield et al., 2002; Terrier et al., 2016). Additionally, behavioral phenomena like context-induced relapse after extinction or punishment that are reliable in rats have not yet been demonstrated in mice (Marchant et al., 2019).

Based on these considerations, we have created and characterized a knock-in rat (*Oprm1*-Cre) that co-expresses the MOR protein and an improved Cre recombinase from the endogenous MOR locus (*Oprm1*). The presence of the Cre transgene did not appear to affect expression of *Oprm1* mRNA or functional MOR protein in the nucleus accumbens (NAc), dorsal striatum (DS), and dorsal hippocampus (dHipp). Similarly, the presence of Cre did not alter MOR-related behaviors in pain-related models or in heroin self-administration and relapse models. These findings show that we generated a knock-in rat for monitoring and manipulating MOR-expressing cells in the rat brain that did not exhibit detectable basal differences in gene expression, protein functionality, or opioid-related behaviors.

We used the knock-in rats to study the role of NAc MOR-expressing cells in heroin self-administration in male and female rats because an early pharmacological study in male rats showed that local NAc injections of the preferential MOR antagonist methyl naloxonium chloride (a lipophobic quaternary derivative of naloxone) decreased the reinforcing effects of self-administered heroin (Vaccarino et al., 1985). We injected the Cre-dependent vector AAV1-EF1a-Flex-taCasp3-TEVP (AAV-DIO-Casp3) (Takahashi et al., 2017) into *Oprm1*-Cre rats and their wildtype littermates, and selectively lesioned NAc MOR-expressing cells in only *Oprm1*-Cre rats. Beyond identifying potential sex differences in the mechanisms of heroin self-administration, our study serves as a proof of concept for the value of this *Oprm1*-Cre rat model in refining our understanding of the functions of endogenous opioid receptor systems.

## Material and Methods

### Subjects

We performed the experiments in accordance with the NIH Guide for the Care and Use of Laboratory Animals (8th edition), under protocols approved by the Animal Care and Use Committees of NIDA Intramural Research Program or the University of Michigan.

#### NIDA

We used 89 *Oprm1*-Cre heterozygotes (41 males and 48 females) and 92 wildtype littermates (45 males and 47 females) for our molecular and behavioral experiments. Before virus or intravenous surgery, the approximate weight range of the rats we used in the behavioral experiments was 350-550 g (males) or 175-300 g (females). We maintained the rats under a reverse 12:12 h light/dark cycle (lights off at 8:00 a.m.) with food and water freely available in the home cage. We housed the rats 2-3 per cage for all experiments except those requiring intravenous surgery which were housed 2-3 per cage prior to surgery and individually after surgery.

#### U Michigan

We used 25 *Oprm1*-Cre heterozygotes (17 males and 8 females) and 25 wildtype littermates (17 males and 8 females) for our molecular and behavioral experiments. We excluded 1 rat due to health problems. The approximate weight range of the rats we used in the behavioral experiments was 350-550 g (males) or 175-300 g (females). We housed the rats 2 per cage (except one rat whose cage mate was excluded due to health problems) in a temperature- and humidity-controlled vivarium, on either a 12-h (lights on at 07:00; males) or a 14-h (lights on at 05:00; females) light cycle, with food and water freely available.

We excluded one female *Oprm1*-Cre rat from the proof-of-concept AAV-DIO-Casp3 experiment due to misplaced injection, one male (Exp. 2) and two female (Exp. 3B & 3C, 1/experiment) *Oprm1*-Cre rats because of poor health, two male rats (Exp. 4, 1/genotype) due to failure to acquire heroin self-administration, and one wildtype female rat (Exp. 5) due to failure to acquire food self-administration. For Exp. 4 & 5, we tested catheters’ patency after the within-dose heroin maintenance phase. We found loss of patency in three *Oprm1*-Cre rats (Exp. 4, 2 males, 1 female) and two wildtype rats (Exp. 5, 1/sex) and only included their data in the food and heroin acquisition phase and food FR phase (Exp. 5 rats).

### Rat *Oprm1* iCre Recombinase Knockin

We used CRISPR/Cas9 technology to introduce the iCre recombinase coding sequence (Shimshek et al., 2002) before the termination codon of the rat *Oprm1* gene. The knockin approach followed the Easi-CRISPR method (Quadros et al., 2017). The canonical rat *Oprm1* gene codes a 2306 nt mRNA (1196 coding bp) from 4 exons that translates a 398 aa protein (NCBI Ref RNA sequence NM_013071.2). We used the CRISPOR algorithm (CRISPR.tefor.net, (Concordet and Haeussler, 2018)) to select a single guide RNA (sgRNA) that was predicted to cut between the second and third base pair of the stop codon located in *Oprm1* exon 4. The guide sequence was 5’ ACTGCTCCATTGCCCTAACT 3’ (PAM = GGG) (Fig. 1). This sgRNA has a high specificity score (CFD = 91, (Doench et al., 2016)).

**Figure 1.**
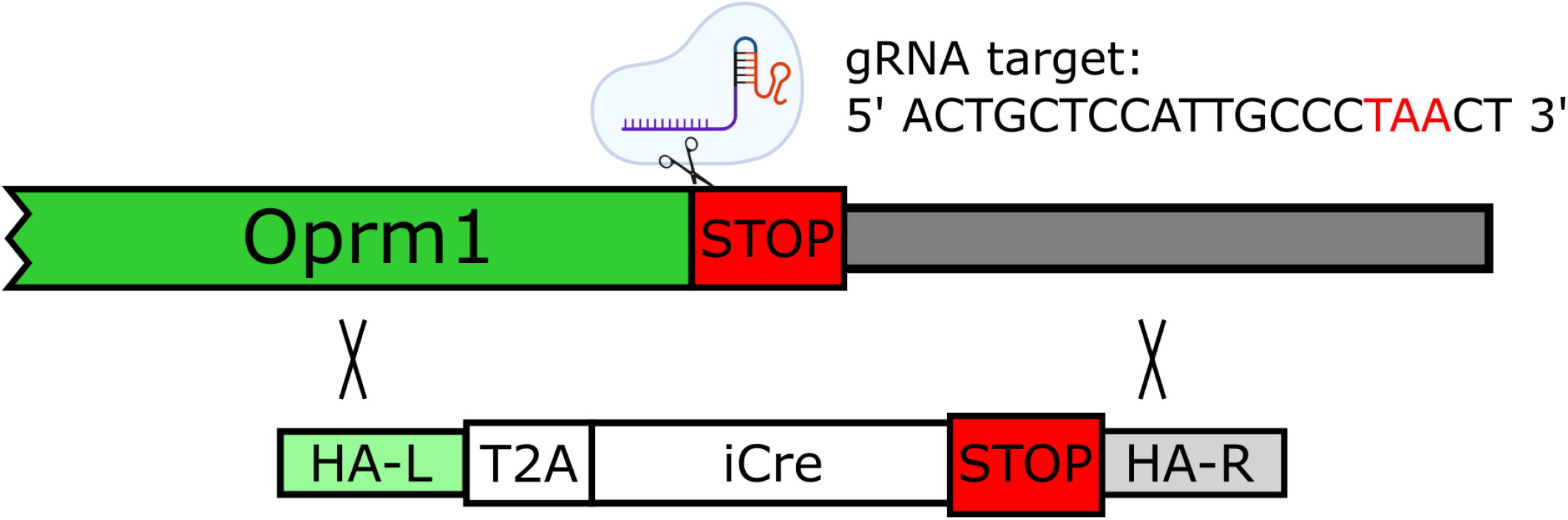
CRISPR-mediated knock-in of T2A-iCre downstream of the rat Oprm1 coding sequence. Schematic of the target gene (rat *Oprm1*) with annotation for the location and sequence of the SpCas9 sgRNA that cleaves within the stop codon. The donor template encoding homologous arms and the T2A-iCre transgene are also shown.

The chemically modified sgRNA (Basila et al., 2017) was synthesized by MilliporeSigma. We tested and verified the sgRNA, which induced Cas9-mediated chromosome breaks, in primary rat embryonic fibroblasts. Briefly, ribonucleoprotein complexes (RNP) were formed by combining 4.85 µg sgRNA with 21.6 µg of enhanced specificity Cas9 protein (eSpCas9, sigmaaldrich.com, (Slaymaker et al., 2016)) in 20 µl. We added RNP to a 0.2 cm cuvette containing 5 µg of a PGKpuro drug resistance plasmid (kind gift of Michael McBurney, (McBurney et al., 1994)) and 750 million cells suspended in D-PBS. A BioRad Gene Pulser was set to deliver a square wave pulse at 250 V, 2 ms pulse width for one pulse with unipolar polarity. We plated cells onto one 6-cm dish in culture medium (high glucose DMEM with the addition of 10% fetal bovine serum, 4 mM glutamine, and Penicillin-Streptomycin at 10,000 U/mL). The next day, we changed the electroporation media. On days 2 and 3, we used media containing 2 µg/µl puromycin to eliminate cells that did not receive the PGKpuro plasmid. On days 4 and 5, we used media without puromycin to feed the cells. On the following day, we collected surviving cells for DNA extraction.

We used PCR primers to amplify a 776 bp genomic DNA fragment that included the sgRNA target (*Oprm1* Activity Forward primer: 5’ ATGGAAGATGGAGCAAGAGAAAGAATTT3’, *Oprm1* Activity Reverse primer: 5’ ATGTATCACTACAGTGAATTTAACAAGTGAC 3’). We submitted amplicons for Sanger sequencing. Chromatograms revealed superimposed peaks, typical of indels formed by non-homologous endjoining (NHEJ) repair of Cas9-induced chromosome breaks (Brinkman et al., 2014).

We used the sgRNA to produce iCre knockin rats because of its high activity. The use of an sgRNA with high specificity in combination with eSpCas9 dramatically reduces the likelihood of Cas9 off target hits in generation zero (G0) founder animals (Anderson et al., 2018). The DNA donor was obtained as a long single stranded DNA Megamer synthesis of 1,312 nucleotides (Table S1A, IDTDNA.com). We designed the DNA donor to include a Gly-Ser-Gly linker and T2A self-cleaving peptide after *Oprm1* codon 398. In this way, *Oprm1* and iCre will be expressed from a single bi-cistronic locus so that physiological levels of MOR protein will be present in cells in addition to the iCre protein (Kim et al., 2011). We performed pronuclear microinjection of 30 mouse zygotes with the RNP and DNA donor to demonstrate that the reagents did not interfere with mouse zygote development *in vitro* with the expectation that such reagents would not then interfere with rat zygote development *in utero*. We obtained mouse zygotes for pronuclear microinjection from B6SJLF1 mice (Jackson Laboratory stock number 100012).

To produce the rat *Oprm1* iCre recombinase knockin, we obtained rat zygotes from Sprague-Dawley rats (Charles River Laboratory Strain Code 001). We performed pronuclear microinjection with a mixture containing RNP (30 ng/µl sgRNA mixed with 50 ng/µl eSpCas9 protein) and 5 ng/µl of the DNA donor as described (Filipiak and Saunders, 2006; Filipiak et al., 2019). Of 125 microinjected rat zygotes, 115 survived microinjection and were surgically transferred to pseudopregnant SAS Sprague-Dawley rats (Charles River Strain Code 400). After 39 possible G0 founder rats were born, we extracted DNA from tail tip biopsies and amplified with primers specific for the iCre coding sequence (Internal iCre Forward primer 5’ AGAAGAAGAGGAAAGTCTCCAACCTGCT 3’ and Internal iCre Reverse primer: 5’ TTTCTGATTCTCCTCATCACCAGGGACA 3’; expected DNA fragment size: 379 bp). After screening 39 potential founder pups, 14 were positive for iCre recombinase.

We screened the 14 iCre positive rats for correct genomic targeting with PCR primers in genomic DNA and in iCre to amplify the 5’ and 3’ junctions of the iCre insertion site. The 5’ junction primers were 5’ Junction Forward primer: 5’ AAGACAATGTTCAGTACAGTTCTCATACC 3’ and 5’ Junction Reverse primer ATTCTCCTTTCTGATTCTCCTCATCAC 3’; expected DNA fragment size from iCre coding sequence insertion: 595 bp. The 3’ junction primers were 3’ Junction Forward primer: 5’ GATGAACTACATCAGAAACCTGGACTC 3’ and 3’ Junction Reverse primer TTCAAGGTGAAAGTTTTAAGTTGGAAATG 3’; expected DNA fragment size from iCre coding sequence insertion: 571 bp. PCR amplicons showed that 1 of the 14 iCre positive rats was positive for both 5’ and 3’ junctions. Sanger sequencing of the amplicons demonstrated iCre was inserted in the desired location.

We also used spanning primers placed in genomic DNA to produce amplicons across the iCre insertion site. Spanning primers were Spanning Forward primer 5’ CAGAGGAAGTCTTCTAACATGCGGTGAC 3’, Spanning Reverse primer 5’ TCTGGATGGTGTGAGACCCAGTTAGTC 3’; expected DNA fragment size: 1123 bp. We gel purified the PCR amplicons and subjected them to TOPO TA cloning and Sanger sequencing to confirm correct iCre insertion into the *Oprm1* gene and that the iCre coding sequence was intact.

We mated the confirmed G0 rat with wildtype Sprague-Dawley rats and obtained germline transmission from the G0 founder. Sequencing of DNA isolated from 14 obligate heterozygote G1 pups showed that they inherited the correctly targeted Oprm1 iCre recombinase knockin. We used G2 rats descended from two (G1) founders for further colony expansion. We housed the rats in an AAALAC accredited facility in accordance with the National Research Council’s guide for the care and use of laboratory animals.

### Breeding and genotyping

#### NIDA and U Michigan

We set up male and female *Oprm1*-Cre heterozygotes in breeding with wildtype mates (CD (Sprague Dawley) IGS, Charles River Labs, strain code #400) for at least 4 generations. At NIDA, we used at least 8 different breeding pairs at each generation after the 3^rd^ generation and used heterozygotes rats and wildtype littermates from the 4^th^ and 5^th^ generation. At U Michigan, we used heterozygotes rats and wildtype littermates from 4 genetically diverse litters from the 7^th^ generation. Tail genotyping was performed by Transnetyx at NIDA and in-house PCR at U Michigan. The rats bred at NIDA were registered with the Rat Genome Database (RG#TBD) and deposited at the Rat Resource and Research Center (RRRC#TBD).

### Fluorescence *in situ* Hybridization Chain Reaction (HCR FISH) (U Michigan)

We designed the Split-initiator DNA probes (*version* 2.0) (Choi et al., 2018; Kumar et al., 2021) (Table S1B and S1C) and the probes were synthesized by Integrated DNA Technologies (Coralville, Iowa, USA). We purchased DNA hairpins conjugated with B3 AlexaFluor-546 (AF-546) and B2 AF-647 from Molecular Instruments (Los Angeles, CA, USA). We sectioned fresh-frozen rat brains at 30 µm in a cryostat: sections were from NAc, DS, and dHipp using AP coordinates from Bregma of +1.2 to +2.0 mm for NAc and DS and -2.4 to -3.0 mm for dHipp. We optimized the HCR FISH method as described previously (Choi et al., 2018; Kumar et al., 2021).

We fixed the sections in 4% paraformaldehyde (PFA), washed with 5X sodium chloride/sodium citrate/0.01% Tween-20 (SSCTw) buffer for 3 times (5 min each), and then acetylated in 0.1M triethanolamine, pH 8.0 with 0.25% vol/vol acetic anhydride solution for 10 min. After rinsing with ddH2O, we de-lipidated the sections in -20°C chilled acetone: methanol (1:1) for 5 min, washed with 5x SSCTw, and equilibrated in hybridization buffer (30% deionized formamide, 5x SSC, 9 mM citric acid [pH 6.0], 0.5 mg/ml yeast tRNA, 1x Denhardt’s solution, 10% dextran sulfate, 0.1% Tween 20) for 60 min, and then incubated the sections in hybridization buffer containing 10 nM initiator-labeled probes at 37°C for 16 h.

After hybridization, we washed the sections at 37°C with probe wash buffer (30% formamide, 5x SSC, 0.1% Tween 20) 3 times and twice with 5x SSCTw for 15 min each. We equilibrated the sections in amplification buffer for 60 min (5x SSC, 10% dextran sulfate, 0.1% Tween 20). We diluted fluorophore-labeled hairpins separately from 3 µM stock to a 2.25 µM final concentration in 20x SSC, heated at 90°C for 90 s, and then snap-cooled to room temperature for 30 min in the dark. We further diluted snap-cooled hairpins to 60 nM final concentration in amplification buffer. We incubated the sections in amplification buffer with hairpins for 16 h at room temperature. Finally, we washed the sections in 5x SSCTw twice for 30 min and mounted and cover-slipped slides with Vectashield antifade mounting medium (Cat #H-1000, Vector Labs).

### Confocal Microscopy (U Michigan)

We acquired image stacks using an Olympus Fluoview-3000 confocal microscope that consisted of 3 channels: DAPI, Cre, and Oprm1. For quantitative colocalization analysis, we used 10x magnification objective lens (Olympus UPLSAPO10X2, numerical aperture (N.A.) 0.4 / working distance (W.D.) 2.2 mm) to acquire image-stacks (xy-dimension 1.59 µm/pixel x 1.59 µm/pixel and z-step of 4.5 µm; 4-6 z-slices per stack). For high magnification representative images, we used 40x (silicone oil immersion, Olympus UPLSAPO40XS, N.A.-1.25/W.D.-0.3 mm) objectives. We selected the image acquisition settings (mainly the PMT Voltage and the laser transmissivity) for optimal pixel saturation to avoid excessive or weak signal and kept these settings constant for all sections.

### Image Processing and Analysis (U Michigan)

We used open-source ImageJ/Fiji software (Schindelin et al., 2012; Schneider et al., 2012) for processing and quantitation, and Amira (Thermo Fisher) for visualization-representation. We processed and quantified images of 2-3 sections per rat (n=5 males/per genotype) per brain area. Regions of interest (ROIs) were drawn following the coronal diagrams and Nissl stain plates (Paxinos and Watson, 2007) and saved in the ROI manager for quantification. 3D image stacks were processed globally, first using the subtract background (rolling=50 stack) tool and then filtered using non-local means denoising method (auto estimate sigma), an adaptive-manifold-based approach which naturally preserves most features of objects and reduces background noise. Next, Gaussian blur (sigma=1) was used to concentrate the signal towards the center of each cell and then segmented using auto local threshold (method=Phansalkar radius=15 parameter_1=0 parameter_2=0). After a watershed split, we estimated *iCre* and *Oprm1* cell numbers in each channel using analyze particle tool (size=15-400, circularity=0.3-1.0). Subsequently, the generated masks of each channel (iCre and Oprm1) were used to quantify the number of colocalized cell bodies through Image Calculator ‘AND’ and Analyze ‘Particle’ tools. We estimated area of ROIs using maximum thresholding values which then were used to quantify neurons per mm^2^ (density number) and percent of colocalized cells and performed statistical analyses on both density and percent data.

### RNAScope *in situ* hybridization and immunohistochemistry (NIDA)

We deeply anesthetized the rats with isoflurane and rapidly decapitated them. We extracted and flash-froze the brains in isopentane solution on dry ice. We sectioned the brains at 20 µm in a cryostat (−13-15°C), air-dried them on the slides at −20°C, and stored them at −80°C. For the detection of *Oprm1* mRNA, we used the RNAscope 2.5 HD Assay – RED kit (322360; Advanced Cell Diagnostics). We fixed the sections for 20 min in neutral buffered 10% formalin, followed by ethanol dehydration series on 50%, 70%, 95%, and 100%, 5 min each. We stored the sections overnight at −20°C in 100% ethanol.

Before hybridization, we incubated the sections in H_2_O_2_ followed by protease IV (322340, Advanced Cell Diagnostics). We hybridized the sections with the *Oprm1* mRNA probe (Cat #410691, Advanced Cell Diagnostics) for 2 h at 40°C and amplified the probe using RNAScope amplifiers as directed by the manufacturer. For brightfield, we detected the *Oprm1* probe with the chromophore Fast Red. We counterstained the sections with methylene blue, removed the excess dye, washed the sections 3 times for 1 min in ddH_2_0, dried the sections in the oven at 60°C for 15 min, cooled the slides to room temperature, dipped them in Citrosolv (Cat #04-355-121, Fisher Scientific), and cover-slipped the slides with Permount (Cat #SP15-100, Fisher Scientific).

### Image processing and analysis (NIDA)

For brightfield microscopy, we imaged *Oprm1* mRNA signal that was labeled Fast Red and nuclei that were labeled with methylene blue. To quantify Capase-3-induced lesions using Fiji ImageJ, we drew the ROI for NAc shell and core on both hemispheres, segmented fast red labeling from methylene blue staining using Giemsa color deconvolution, made thresholds for red grains, and segmented from channel 2 of the color deconvolution. We divided the pixels covered by grains by the pixels within the ROI to determine the percent of area covered by the *Oprm1* mRNA signal (% area covered by red grains).

### [^35^S]GTPγS autoradiography

We cut frozen brain sections at 20 µm using a cryostat and thaw mounted the sections onto glass slides. We pipetted pre-incubation buffer onto each slide and incubated for 20 min at room temperature (50 mM Tris-HCl, 1 mM EDTA, 5 mM MgCl_2_ and 100 mM NaCl). We removed the buffer by aspiration and incubated the sections for 60 min in pre-incubation buffer containing 2 mM GDP and 1 µM DPCPX. We removed the GDP buffer and pipetted [^35^S]GTPγS cocktail (GDP buffer, 1 mM DTT, 0.625 nM [^35^S]GTPγS) with DAMGO (10 μM), without DAMGO (basal condition), or with a saturated concentration of non-radioactive GTP (for non-specific binding) onto each slide and incubated for 90 min. We washed (2 × 5 min, 50 mM Tris-HCl, pH 7.4, 5 mM MgCl_2_), rinsed (30 sec in ice water), and air-dried the slides. Along with radioactive standards (nCi/g), apposed them to a BAS-SR2040 phosphor screen (Fujifilm) for 3 days and imaged the slides using a phosphorimager (Typhoon FLA 7000; GE Healthcare). We drew ROIs onto the sections with standards using Multigauge software (GE Healthcare) and expressed values as % basal.

### [^3^H]DAMGO autoradiography

We cut frozen brain sections at 20 µm using a cryostat and thaw mounted the sections onto SuperPlus glass slides (Avantor). We pre-incubated slides (10 min, room temperature) in incubation buffer (50 mM Tris-HCl, pH 7.4), then incubated (60 min, room temperature) in incubation buffer containing [^3^H]DAMGO (5 nM). We determined non-specific binding in the presence of 10 µM naloxone. After incubation, we washed (2 × 30sec, incubation buffer), rinsed (30sec in ice water), and air dried the slides and along with radioactive standards (nCi/g), apposed them to a BAS-TR2025 Phosphor Screen (Fujifilm) for 10 days and imaged using a phosphorimager (Typhoon FLA 7000). We drew ROIs onto the sections with standards using Multigauge software (GE Healthcare) and expressed and calibrated values as nCi/g.

### Apparatus (food and drug self-administration)

We trained and tested the rats in standard Med Associates self-administration chambers. Each chamber had two retractable levers located 7.5-8 cm above the grid floor on the right wall with a food receptacle between them, and an inactive non-retractable lever on the left side. A tone cue is located above one of the levers and a light cue is located above the other lever. Lever presses on the retractable levers activated either the infusion pump or a pellet dispenser. Lever presses on the inactive lever had no programmed consequences. In Experiment 2, the self-administration and extinction contexts differed in their auditory, visual, and tactile cues, as in our previous studies (Adhikary et al., 2017; Bossert et al., 2019). We refer to the contexts as A and B, where A is the context of self-administration training and reacquisition, and B is the context of extinction. We counterbalanced the physical environments of contexts A and B.

### Drugs

#### NIDA

We received heroin hydrochloride (HCl) and morphine sulfate from the NIDA pharmacy and dissolved them in sterile saline. The heroin unit doses of Experiment 2 are based on our previous work (Bossert et al., 2004; Bossert et al., 2016; Bossert et al., 2022) and the heroin unit doses in Experiment 4-5 are based on Stewart et al. (1996). The morphine (1 ml/kg, s.c) doses and lactic acid (Cat #L1250, Sigma Aldrich, dissolved in sterile water) concentrations in Experiment 3 are based on Baldwin et al. (Baldwin et al., 2022) and Reiner et al. (Reiner et al., 2021). We injected morphine 30 min prior to behavioral testing or 20 min prior to lactic acid injections; we injected lactic acid 10 min prior to behavioral testing.

### U Michigan

We prepared morphine sulfate from a pharmaceutical liquid stock (Mitigo USP preservative free, 25 mg/mL, Piramal Critical Care) that we diluted with sterile saline (Fresenius Kabi). We dissolved naloxone HCl (Tocris Bioscience) in sterile saline. We injected both drugs i.p. in a volume of 1 ml/kg. We injected naloxone 30 min prior to morphine, which we injected 60 min prior to behavioral testing.

### Surgery

#### Intracranial surgery for viral delivery

In our proof-of-concept experiment, we injected AAV1-EF1a-Flex-taCasp3-TEVP (NIDA Genetics & Engineering Viral Vectors Core (GEVVC), lot # AAV-2015-11-10-B, titer: 5.16E+11 vg/ml) unilaterally into the right hemisphere of NAc shell and phosphate-buffered saline into the left hemisphere; injections were 500 nl/side. In Experiment 4, we injected AAV1-EF1a-Flex-taCasp3-TEVP (NIDA GEVVC, lot # AAV-2015-11-10-B, titer: 5.16E+11 vg/ml) bilaterally into NAc shell; injections were 1000 nl/side. In Experiment 5, we injected AAV1-EF1a-DIO-EYFP (NIDA GEVVC, lot # AAV-2015-02-17-B, titer: 2.73E+12 vg/ml) bilaterally into NAc shell; injections were 1000 nl/side. We used the following coordinates from Bregma: AP, 1.6 mm; ML, 2.5 mm (10° angle); DV, -7.5 mm (males) and - 7.3 mm (females). These coordinates are based on a previous study (Marchant et al., 2016). We delivered the AAVs using Nanofil syringes (WPI, 33 gauge) at a rate of 100 nl/min. After each injection, we left the injection needle in place for 3 min to allow for diffusion. After the injections, we filled the drilled holes with bone wax and closed the wounds using autoclips (Texas Scientific Instruments).

#### Intravenous surgery

We anesthetized the rats with isoflurane (5% induction; 2-3% maintenance, Covetrus). We attached silastic catheters to a modified 22-gauge cannula cemented to polypropylene mesh (Industrial Netting), inserted the catheter into the jugular vein, and fixed the mesh to the mid-scapular region of the rat (Caprioli et al., 2015; Fredriksson et al., 2020). We injected the rats with ketoprofen (2.5 mg/kg, s.c., Covetrus) during surgery and on the following day to relieve pain and decrease inflammation. We also injected Enrofloxacin (2.27% diluted 1:9 in sterile saline, s.c., Covetrus) during surgery, 4-5 days post-surgery, and if we observed an infection during the experiment. The rats recovered for 6-8 days before heroin self-administration training. During all experimental phases, we flushed the catheters daily with gentamicin in sterile saline (4.25 mg/ml, 0.1 ml, Fresenius Kabi).

### Behavioral experiments

#### Experiment 1: Food self-administration

The goal of Experiment 1 was to determine if there are differences between *Oprm1*-Cre rats and their wildtype littermates in operant learning and performance for a non-drug reward. For this purpose, we used 45 mg high carbohydrate food pellets (TestDiet, Cat # 1811155) that food-sated rats strongly prefer over heroin, fentanyl, and methamphetamine (Caprioli et al., 2015; Venniro et al., 2017; Reiner et al., 2020). We trained and tested the rats (8 males, 12 females) approximately 5 h after the onset of the dark cycle (8 am). The experiment consisted of two phases: (1) acquisition of food self-administration for 7 days for 1-h/d under a fixed-ratio 1 (FR1) 20-s timeout reinforcement schedule, and (2) tests for food self-administration after increasing the response requirements from FR1 to FR8 in the following sequence: FR1 (3 d), FR2 (1 d), FR4 (1 d), FR6 (1 d), and FR8 (1 d).

Each session began with the illumination of the houselight and the insertion of the food-paired active lever 10 s later. During the first 7 daily acquisition sessions, we mildly food restricted the rats (removed their food between 8 and 9 am) and gave them 1-h magazine-training sessions before the operant training during which 1 pellet was delivered noncontingently every 2 min. Lever presses led to the delivery of one 45-mg pellet and each pellet delivery was paired with a 20-s white-light cue.

#### Experiment 2: Heroin self-administration and relapse-related behaviors

The goal of Experiment 2 was to determine if there are differences between *Oprm1*-Cre rats and their wildtype littermates in heroin self-administration and commonly used relapse-related behaviors: extinction responding, context-induced reinstatement, and reacquisition (Bossert et al., 2013; Venniro et al., 2016; Khoo et al., 2017). We used a variation of the ABA context-induced reinstatement (renewal) procedure in which rats are trained to self-administration heroin in context A, are tested for extinction of heroin-reinforced responding in context B, and then tested for context-induced reinstatement of heroin seeking and reacquisition in context A (Bossert et al., 2020; Bossert et al., 2022).

##### Training in context A (12 days)

We trained the rats (12 males, 16 females) to self-administer heroin HCl in context A for 6 h/day (six 1-h sessions separated by 10 min) for 12 days. Each session began with the illumination of the houselight that remained on for the entire session; the active lever was inserted into the chamber 10 s after the houselight was illuminated. During training, the rats earned heroin infusions by pressing on the active lever; infusions were paired with a compound tone–light cue for 3.5 s under an FR1 20-s timeout reinforcement schedule. Heroin was infused at a volume of 100 µl over 3.5 s at a dose of 100 µg/kg/infusion (first 6 sessions) and then 50 µg/kg/infusion (last 6 sessions). Lever presses on the active lever during the timeout period were recorded but did not result in heroin infusions. Presses on the inactive lever were recorded but had no programmed consequences. At the end of each 1-h session, the houselight turned off and the active lever was retracted. If we suspected catheter failure during training, we tested patency with Diprivan (propofol, NIDA pharmacy, 10 mg/mL, 0.1-0.2 mL injection volume, i.v.).

##### Extinction responding in context B (7 days)

We ran the rats under extinction conditions in context B for 6-h per day (six 1-h sessions separated by 10 min) for 7 days. During this phase, presses on the previously active lever led to presentation of the discrete tone-light cue but not heroin infusions.

##### Context-induced reinstatement in contexts A and B (2 d)

We tested the rats under extinction conditions (see above) for 6-h per day for 2 days in context A and context B in a counterbalanced order.

##### Reacquisition of heroin self-administration in context A (1 d)

We tested reacquisition of heroin self-administration during one 6-h session in context A. During testing, lever presses were reinforced by heroin (50 µg/kg/infusion, FR1 20-s timeout reinforcement schedule) and the discrete tone-light cue. After the 6-h session, we tested catheter patency with propofol (NIDA pharmacy, 10 mg/mL, 0.1-0.2 mL injection volume, i.v.).

#### Experiment 3: Evaluation of pain-related responses using von Frey test, tail flick test, and lactic acid-induced suppression of operant responding

The goal of Experiment 3 was to determine if there are differences between *Oprm1*-Cre rats and their wildtype littermates in pain sensitivity and morphine analgesia using three different methods: von Frey test, tail flick test, and lactic acid-induced suppression of operant responding.

##### Experiment 3a: Von Frey test

We performed all testing in a quiet, dimly lit room and we gave the rats about 30 min to habituate to the testing environment before testing began. We assessed sensitivity to mechanical stimulation using nylon von Frey filaments (BiosEB). We placed the rats (14 males, 6 females) on a stainless steel 1 cm square mesh grid and applied von Frey filaments to the plantar surface of both hind paws using the sampling method described in (Wang et al., 2005). We obtained paw withdrawal threshold scores for both hind paws and averaged them to produce a single composite withdrawal score for each rat. Following an initial test of baseline response, we injected rats with increasing doses of morphine (0.625, 1.25, and 2.5 mg/kg, i.p.) 60 min prior to test, 1 dose per day. After acute analgesia testing, we injected naloxone (1 mg/kg, i.p.) 30 min prior to morphine (2.5 mg/kg), which was injected 60 min prior to test. Following this test, we injected the rats daily for 21 days with 2.5 mg/kg morphine to induce analgesic tolerance, and then retested their analgesic response to this dose of morphine. We compared the analgesic response after tolerance development to the response to the same dose during the acute analgesic phase of the experiment.

##### Experiment 3b: Tail flick test

We performed all testing in a quiet, dimly lit room and we gave the rats about 30 min to habituate to the testing environment before testing began. We used a commercially available tail flick apparatus (IITC Life Science) to assess latency to tail flick from a noxious thermal stimulus. We calibrated the testing intensity to provide reliable tail flick latencies of about 4 s at baseline, and an automatic cutoff time of 15 s to prevent tissue damage. We gently restrained the rats (10 males, 10 females) and then applied the noxious thermal stimulus to the caudal 1/3 of the tail and automatically recorded latency to flick the tail away from the stimulus. We conducted 2 tests per rat at least 1 min apart at distinctive locations along the tail to prevent sensitization. We averaged the latencies to produce a single composite latency score for each rat. After an initial test of baseline response, we injected the rats with increasing doses of morphine (1.25, 2.5, 5, and 10 mg/kg, i.p.) 60 min prior to test, 1 dose per day. After acute analgesia testing, we injected naloxone (1 mg/kg, i.p.) 30 min prior to morphine (5 mg/kg), which was injected 60 min prior to test. Following this test, we injected the rats daily for 21 days with 5 mg/kg morphine to induce analgesic tolerance, and then retested their analgesic response to this dose of morphine. We compared the analgesic response after tolerance development to the response to the same dose during the acute analgesic phase of the experiment.

##### Experiment 3c: Lactic acid-induced behavioral depression

Using the 45 mg pellets described in Experiment 1, we trained and tested the rats (8 males, 8 females) approximately 5 h after the onset of the dark cycle (8 am). The experiment consisted of four phases: (1) acquisition of food self-administration for 6 days for 1-h/d under a fixed-ratio 1 (FR1) 20-s timeout reinforcement schedule, (2) acute injections of lactic acid (0, 0.9%, 1.3%, and 1.8%) 10 min prior to food self-administration sessions, (3) acute injections of morphine (0, 1, 3, and 10 mg/kg) 30 min prior to food self-administration sessions, and (4) acute injections of morphine (0, 1, and 3 mg/kg) 20 min prior to acute injections of lactic acid (1.8%) 10 min prior to food self-administration sessions. All injections were counterbalanced within each phase, and we ran baseline sessions (no injections) in between every test day during all phases. During the first 3 acquisition sessions, we mildly food restricted the rats (removed their food between 8 and 9 am) and gave them 1-h magazine-training sessions before the operant training as described above. Lever presses led to the delivery of one 45-mg pellet and each pellet delivery was paired with a 20-s white-light cue.

#### Experiment 4: Effect of Cre-dependent AAV1-EF1a-Flex-taCasp3-TEVP (AAV-DIO-Casp3) NAc lesions on acquisition and maintenance of heroin self-administration

MORs in NAc core and shell are critical for the operant reinforcing effects of heroin in male rats (Vaccarino et al., 1985; Wise, 1989; Koob, 1992). Based on this knowledge, the goal of Experiment 4 was to behaviorally validate the *Oprm1*-Cre knock-in rat by demonstrating that Cre-dependent lesions of NAc MOR-expressing neurons will decrease acquisition and maintenance of heroin self-administration in *Oprm1*-Cre rats.

Experiment 4 consisted of five phases: 1) concurrent acquisition of food (morning) and heroin (afternoon) self-administration (12 days, 3-h/d, 3 days/heroin dose), 2) within-session heroin dose-response (1 day, 2-h/dose), 3) within-session heroin FR response (1 day, 1-h/FR requirement), 4) extended access heroin session (1 day, 9 h), and 5) within-session food FR response (1 day, 1-h/FR requirement).

##### Acquisition: Food and heroin self-administration

We trained the rats (28 males, 28 females) to self-administer food and heroin with a 3-h food session (1 pellet per reward delivery) in the morning and a 3-h heroin session in the afternoon; food pellets were paired with a 20-s white-light cue on one lever and heroin infusions were paired with a 5-s tone on a different lever. Heroin was infused at a volume of 100 µl over 3.5 s at a dose of 12.5 µg/kg/infusion (the first 3 days) and then 25, 50, and 100 µg/kg/infusion for each subsequent 3 days. This acquisition procedure is based on a previous study of Stewart et al. (1996).

##### Heroin maintenance: Within-session dose response

After the rats learned to self-administer heroin, we tested them using an ascending within-session dose-response curve procedure (Deroche et al., 1999; Fredriksson et al., 2017). We tested the ascending heroin doses of 12.5, 25, 50, 100 µg/kg/infusion for 2-h per dose under an FR1 20-s timeout reinforcement schedule. We used two sets of stock solutions and manipulated the intended drug dose by using 2 different infusion times (1.75 s for the 12.5 and 50 µg/kg unit doses, and 3.5 s for the 25 and 100 µg/kg unit doses). We excluded 2 male rats (n=1 per genotype) because they showed no evidence of acquiring heroin self-administration (3 infusions per day or less during the 12 acquisition sessions) despite having patent catheters. We also excluded 3 *Oprm1*-Cre rats (2 males, 1 female) because of loss of catheter patency when tested at the end of the within-session dose response. We also excluded these rats from the subsequent tests described below.

##### Heroin maintenance: Ascending within-session fixed-ratio response

We tested the rats in 1-h consecutive sessions for heroin self-administration (50 µg /kg/infusion) under FR1, FR2, FR4, FR8, FR16, FR32, FR64 reinforcement schedules. This procedure is based on a study of Chow et al. (Chow et al., 2022).

##### Heroin maintenance: Extended access

We tested the rats in a single 9-h heroin (50 µg/kg/infusion, FR1 20-s timeout reinforcement schedule) self-administration session. We retested one rat whose tubing was disconnected during the test session for determining heroin FR response.

##### Food self-administration: Ascending within-session FR response

We tested some of the rats (15 males, 11 females) in 1-h consecutive sessions for food self-administration under FR1, FR2, FR4, FR8, FR16, FR32, FR64 reinforcement schedules. We ran Experiment 4 in two cohorts, and only performed this test on the second cohort.

#### Experiment 5: Effect of control virus AAV1-EF1a-DIO-EYFP (AAV-DIO-EYFP) into NAc on acquisition and maintenance of heroin self-administration

The goal of Experiment 5 was to demonstrate the specificity of the effect of AAV-DIO-Casp3 in *Oprm1*-Cre rats by using a control virus. For this purpose, we injected a virus that has the same Cre-dependent mechanism and same promotor as AAV-DIO-Casp3 but does contain the taCasp-TEVP component that activates the cells and induces apoptosis.

Experiment 5 consisted of the same five phases as Experiment 4 and was run in the same sequence (13 males, 15 females).

### Statistical analysis

We analyzed datasets without any missing values with GLM procedure of SPSS (version 27). We analyzed datasets with missing values with linear mixed effects modeling (Gelman and Hill, 2006) in JMP 16. Specifically, for the von Frey and tail flick tests, we analyzed the dose-response data with Genotype (nominal) as a fixed between-subjects factor, Dose (nominal) as a fixed within-subjects factor, and Subject as a random factor. For the Morphine + Naloxone and Morphine + Tolerance data, we analyzed the data with Genotype (nominal) as a fixed between-subjects factor, Treatment Condition (nominal) as a fixed within-subjects factor, and Subject as a random factor. We also used lined mixed effect modelling in JMP 16 to analyze the heroin self-administration data because some rats were disconnected from the tubing line (9 events out of 660 events for acquisition and 3 events out of 450 events for extended access). For acquisition, we analyzed the data with Genotype and Sex (both nominal) as fixed between-subjects factors, Dose (nominal) as a fixed within-subjects factor, and Subject as a random factor. For extended access, we analyzed the data with Genotype and Sex (both nominal) as fixed between-subjects factors, Hour (nominal) as a fixed within-subjects factor, and Subject as a random factor.

For complete statistical results, see Supplemental Results section and Supplemental Table 2 (Table S2). In the Figures, we indicate post-hoc (Fisher PLSD test) genotype differences between each sex and within each sex after significant main effects or interactions (see Results). Because our ANOVAs yielded multiple main and interaction effects, we only report statistical effects that are critical for data interpretation in the Results section. We only used Sex as an experimental factor in Experiment 4 because it is the only experiment that was statistically powered to detect sex differences.

## Results

### Anatomical and cellular validation of the CRISPR-mediated knock-in of T2A-iCre

Fig. 1 shows a schematic of the target gene (rat *Oprm1*) with annotation for the location and sequence of the SpCas9 gRNA that cleaves immediately before the stop codon.

#### HCR FISH assay

We used HCR FISH to label *Oprm1, iCre*, and *Oprm1+iCre+* mRNA double-labeled cells in *Oprm1*-Cre male rats and their wildtype littermates. We found no genotype differences in *Oprm1+* cells per mm^2^ in NAc, DS, or dHipp (Fig. 2A). We found a significant number of double-labeled *Oprm1+/Cre+* double positive labeled cells (compared to *Oprm1+Cre* negative cells) in *Oprm1*-Cre rats in all brain areas: NAc: F(1,4)=356.1, p<0.001, DS: F(1,4)=652.4, p<0.001, dHipp: F(1,4)=172.3, p<0.001 (Fig. 2B). The percent of *Cre+/Oprm1*+ cells in the different brain areas was 95-98% (Fig. 2C) and Cre was not detected in wildtype littermates.

**Figure 2.**
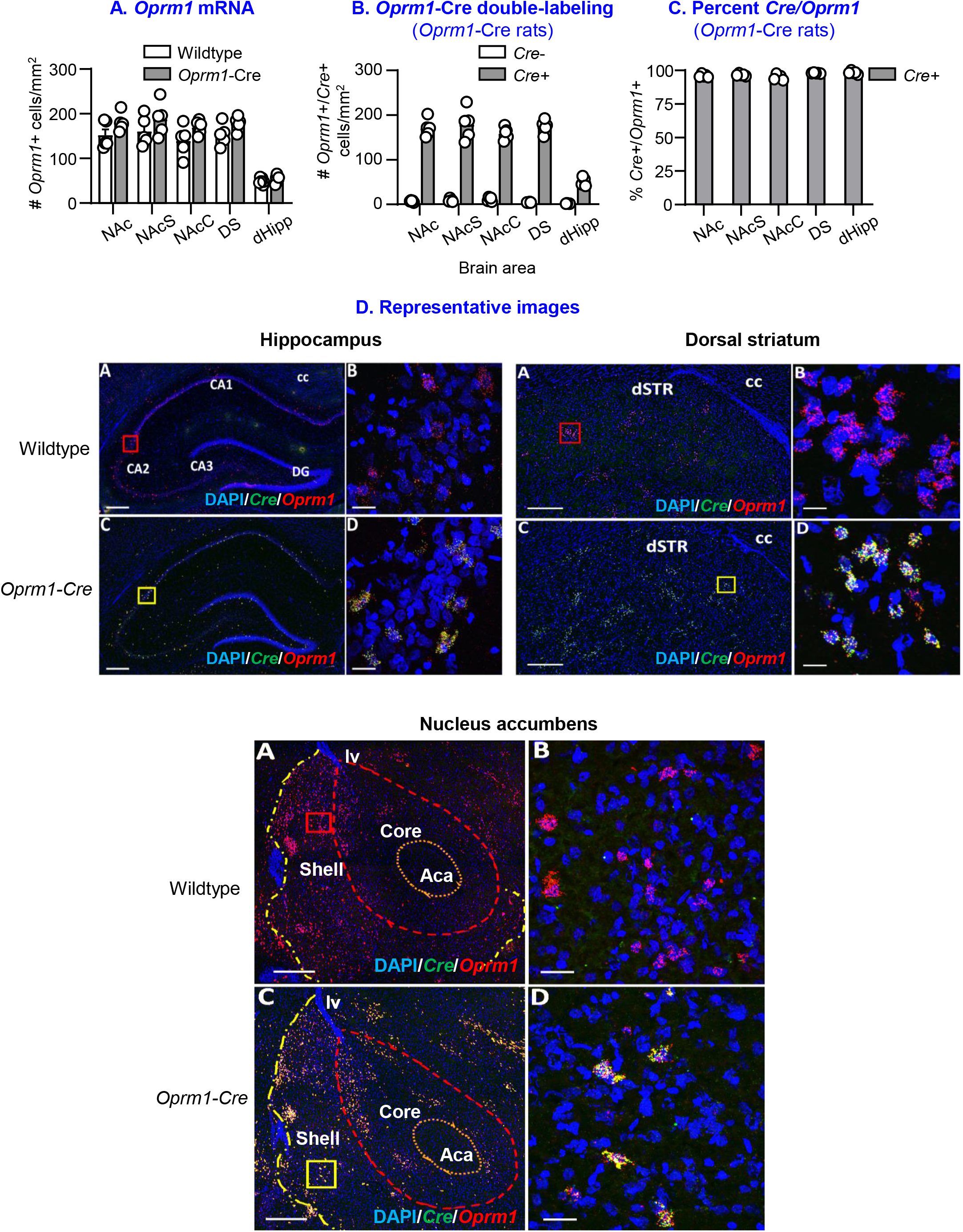
iCre mRNA and Oprm1 mRNA in nucleus accumbens, dorsal striatum, and dorsal hippocampus. **(A)** *Oprm1+* cells per mm^2^ for *Oprm1* mRNA (wildtype and *Oprm1*-Cre, n=5/genotype). **(B)** *Oprm1+*/*Cre+* double labeled cells per mm^2^ (*Oprm1*-Cre rats only). **(C)** Percent *Cre+/Oprm1+* (*Oprm1*-Cre rats only). **(D)** Representative confocal photomicrographs of *Oprm1-*Cre rat brains showing colocalization (**yellow**) between *Oprm1*^*+*^ neurons (**red**) and *Cre*^*+*^ neurons (**green**) in dHipp, DS, and NAc in comparison to wildtype rats which only showed *Oprm1* expression (**red**). Objective lens magnification: **D: A**,**C** 10X and **D: B**,**D** 40X. Scale bars: **D: A**,**C** = 300 µm; **D: B**,**D** = 25 µm. <u>Abbreviations:</u> anterior commissure, Aca; hippocampal subfields, CA1, CA2, CA3; corpus callosum, cc; dentate gyrus, DG; dorsal striatum, dSTR; left ventricle, lv.

#### [^35^S]GTPγS autoradiography and [^3^H]DAMGO autoradiography assays

We used autoradiography to measure MOR activity (via [^35^S]GTPγS) and binding (via [^3^H]DAMGO) in *Oprm1-*Cre male and female rats and their wildtype littermates. We found no genotype differences in DAMGO-stimulated [^35^S]GTP*γ*S recruitment in NAc and DS (p>0.05) (Fig. 3A). We also found no genotype differences in [^3^H]DAMGO binding in NAc (p>0.05). However, [^3^H]DAMGO binding was higher binding in DS in *Oprm1*-Cre rats than in wildtype littermates (F(1,10)=6.7, p=0.027) (Fig. 3B).

**Figure 3.**
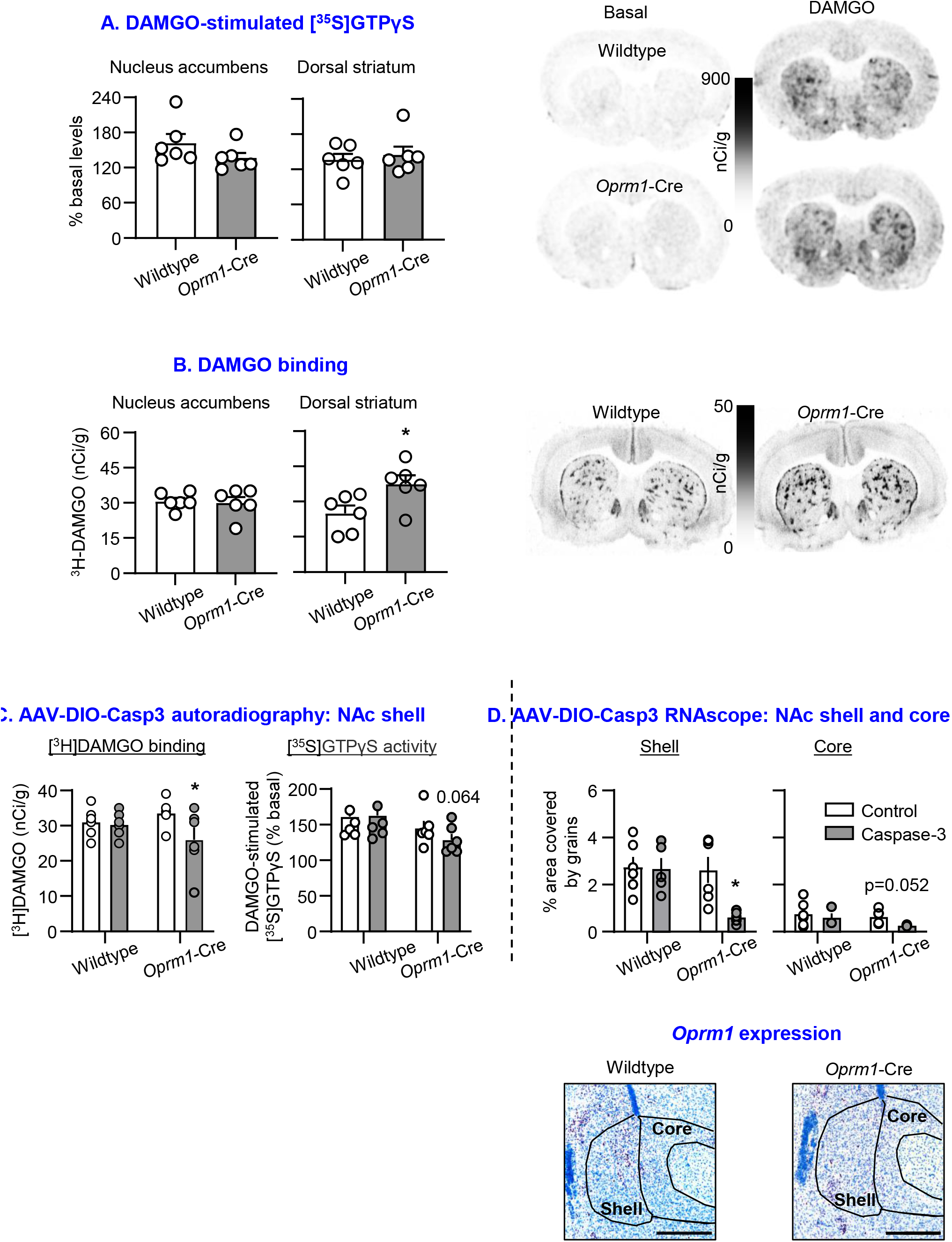
Autoradiography in Oprm1-Cre rats and wildtype littermates. **(A)** DAMGO-stimulated [^35^S]GTPyS in NAc and DS in wildtype and *Oprm1-*Cre rats (n=6/genotype & sex). Values are calibrated and expressed as % basal. **(B)** [^3^H]DAMGO binding in NAc and DS in wildtype and *Oprm1-*Cre rats (n=6/genotype & sex). Values are calibrated and expressed as nCi/g. *AAV-DIO-Casp3 lesion in nucleus accumbens: autoradiography and RNAscope*. We injected AAV1-EF1a-Flex-taCasp3-TEVP unilaterally into the right hemisphere of NAc shell and phosphate-buffered saline into the left hemisphere; injections were 500 nl/side. **(C)** [^3^H]DAMGO binding (left panel) and DAMGO-stimulated [^35^S]GTPyS (right panel) in NAc of wildtype and *Oprm1-*Cre rats (n=5-6/sex & genotype). Values are calibrated and expressed as nCi/g and % basal, respectively. **(D)** Mean±sem *Oprm1*+ cells expressed as red grains per area (% area covered by red grains) in NAc shell and core in wildtype and *Oprm1-*Cre rats (n=5-6/genotype & sex). Scale bar = 500 µm. * Different from the control hemisphere, p<0.05.

#### AAV-DIO-Casp3 lesion in NAc: [^3^H]DAMGO autoradiography and RNAscope

We injected AAV-DIO-Casp3 unilaterally (500 nl/side) into NAc shell and measured *Oprm1* mRNA expression and MOR activity and binding in *Oprm1-*Cre male and female rats and their wildtype littermates. The analysis of DAMGO binding and DAMGO-stimulated [^35^S]GTPγS recruitment in NAc, which included the between-subjects factor of Genotype (Wildtype, *Oprm1*-Cre) and the within-subjects factor of Lesion (Vehicle, AAV-DIO-Casp3), showed significant Genotype x Lesion interaction for binding (F(1,9)=8.4, p=0.018) and approaching significant interaction for DAMGO-stimulated [^35^S]GTPγS activity (F(1,9) = 4.8, p=0.056) (Fig. 3C). The analysis of % area covered by red grains in NAc shell and core, which included the between-subjects factor of Genotype and the within-subjects factor of Lesion, showed significant effects of Genotype x Lesion interaction in NAc shell (F(1,9)=9.9, p=0.012), but not in NAc core. (Fig. 3D).

Together, these results indicate that AAV-DIO-Casp3 NAc shell injections selectively decreased *Oprm1* mRNA and MOR binding activity in the injected hemisphere of *Oprm1*-Cre male and female rats, but not in the vehicle-injected hemisphere or in wildtype littermates.

### Experiment 1: Food self-administration

There were no genotype differences in acquisition of food self-administration and subsequent responding under the different FR requirements (Fig. 4A). The analysis of acquisition data, which included the between-subjects factor of Genotype (Wildtype, *Oprm1*-Cre) and the within-subjects factor of Session (1 to 7) showed a significant effect of session for both the number of pellets and number of active lever presses (F(6,108)=24.1, p<0.001; F(6,108)=7.8, p<0.001) but no significant effects of Genotype or interaction between the two factors (p>0.1). The analysis of the FR response data, which included the between-subjects factor of Genotype and the within-subjects factor of FR requirement (FR1 to FR8), showed a significant effect of FR requirement for both the number of pellets and number of active lever presses (F(4,72)=34.4, p<0.001; F(4,72)=12.3, p<0.001) but no significant effects of Genotype or interaction between the two factors (p>0.1).

**Figure 4.**
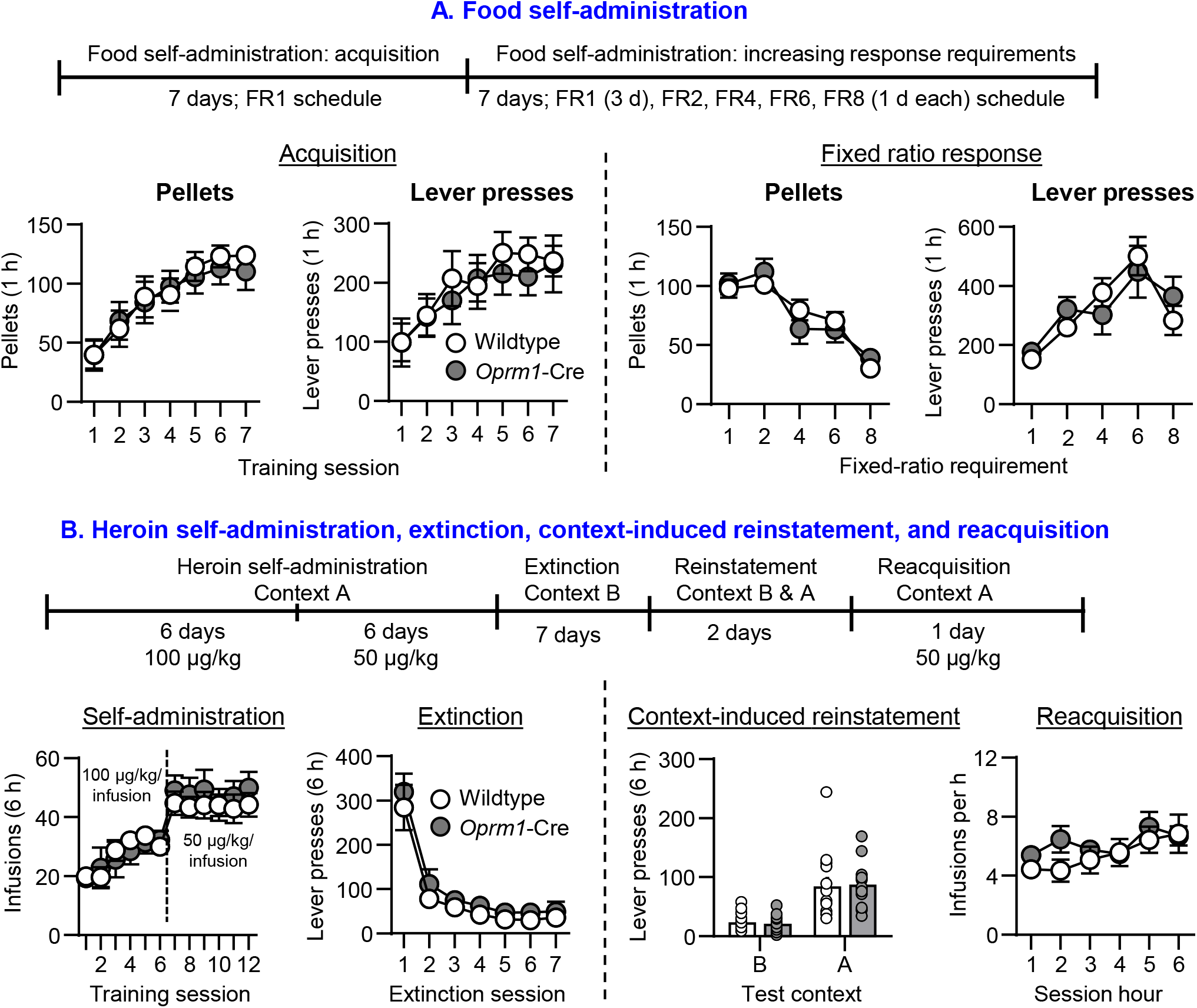
Food self-administration, heroin self-administration, and heroin relapse-related behaviors in Oprm1-Cre rats and wild-type littermates. **(A)** <u>Food self-administration: Acquisition (Left) and Fixed ratio response (Right):</u> Mean±sem number of pellets consumed (Left panel) and active lever presses (Right panel). Wildtype (3 males, 6 females), *Oprm1*-Cre (4 males, 6 females); data were combined for males and females. **(B)** <u>Heroin self-administration:</u> Mean±sem number of heroin infusions during heroin self-administration training (days 1-6, 0.1 mg/kg/infusion; days 7-12, 0.05 mg/kg/infusion). <u>Extinction responding</u>: Mean±sem number of active lever presses during the seven 6-h extinction sessions. Active lever presses led to contingent presentations of the tone-light cue, but not heroin. <u>Context-induced reinstatement:</u> Mean±sem number of active lever presses during the 6-h reinstatement tests in context B and context A. Active lever presses led to contingent presentations of the tone-light cue, but not heroin. <u>Reacquisition:</u> Mean±sem number of heroin infusions (0.05 mg/kg/infusion) per hour during reacquisition. Active lever presses led to the delivery of heroin infusions and the tone-light cue. Wildtype (6 males, 8 females), *Oprm1*-Cre (5 males, 8 females); data were combined for males and females.

Together, the results of Experiment 1 indicate that the knock-in manipulation had no effect on acquisition of palatable food self-administration in mildly food-restricted rats or the effort to self-administer food pellets in food-sated rats.

### Experiment 2: Heroin self-administration and relapse-related behaviors

#### Heroin self-administration (context A)

There were no genotype differences in acquisition of heroin self-administration (Fig. 4B, far left). The statistical analysis of number of infusions, which included the between-subjects factor of Genotype, and the within-subjects factors of Training session (1 to 7) and Training dose (50, 100 µg/kg/infusion), showed significant effects of Training session x Training dose (F(5,125)=10.0, p<0.001). There were no significant effects of Genotype or interactions with this factor (p>c0.1).

#### Extinction responding (context B)

There were no genotype differences in extinction of heroin self-administration (Fig. 4B, mid left). The statistical analysis of number of active lever presses, which included the between-subjects factor of Genotype and the within-subjects factor of Extinction session, showed a significant effect of Extinction session (F(6,150)=49.9, p<0.001) but no significant effects of Genotype or interaction between the two factors (p>0.1).

#### Context-induced reinstatement (contexts A & B)

There were no genotype differences in context-induced reinstatement of heroin seeking (Fig. 4B, mid right). The statistical analysis of number of active lever presses, which included the between-subjects factor of Genotype and the within-subjects factor of Context (A, B), showed a significant effect of Context (F(1,25)=65.4, p<0.001) but no significant effects of Genotype and interaction between the two factors (p>0.1).

#### Reacquisition (context B)

There were no genotype differences in reacquisition of heroin self-administration (Fig. 4B, far right). The statistical analysis of number of infusions, which included the between-subjects factor of Genotype and the within-subjects factor of Session hour (1 to 6), showed significant effects of Session hour (F(5,125)=6.0, p<0.001) but no significant effects of Genotype or interaction between the two factors (p>0.1).

Together, the results of Experiment 2 indicate that the knock-in manipulation had no effect on heroin self-administration, extinction responding, context-induced reinstatement, and reacquisition of heroin self-administration.

### Experiment 3: Evaluation of pain-related responses using von Frey test, tail flick test, and lactic acid-induced suppression of operant responding

#### Experiment 3a: von Frey test

There were no genotype differences in the morphine dose-response of paw withdrawal thresholds (Fig. 5A, left). The statistical analysis of threshold, which included the between-subjects factor of Genotype and the within-subjects factor of Dose (0, 0.625, 1.25, 2.5 mg/kg), showed significant effects of Dose (F(3,47.72)=73.0, p<0.001) but no significant effects of Genotype or interaction between the two factors (p>0.1). There were also no genotype differences in response to naloxone or to the analgesic tolerance to morphine (Fig. 5A, middle and right). The statistical analysis of threshold, which included the between-subjects factor of Genotype and the within-subjects factor of Condition (Morphine + Naloxone, or Morphine + Tolerance), showed significant effects of Condition for response to naloxone (F(2,17.45)=65.4, p<0.001) and for analgesic tolerance (F(2,30.54)=87.4, p<0.001), but no significant effects of Genotype or interaction between the two factors (p>0.1).

**Figure 5.**
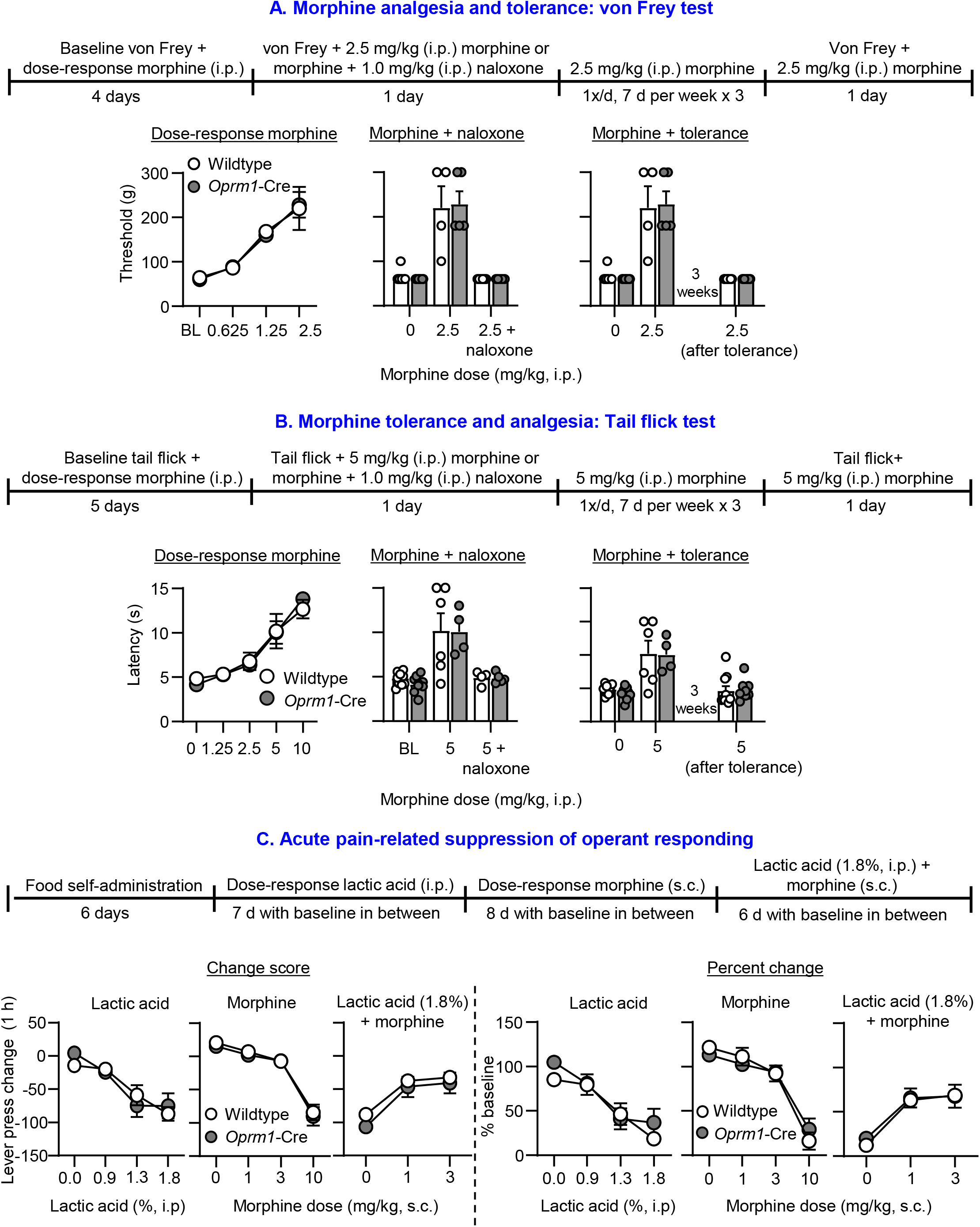
Morphine analgesia, tolerance, and pain-related suppression in Oprm1-Cre rats and wild-type littermates. **(A)** <u>von Frey test and timeline of experiment for morphine analgesia and tolerance</u>. <u>Left panel:</u> Baseline and von Frey thresholds (g) after ascending doses of morphine (0.625, 1.25, and 2.5 mg/kg, i.p.). <u>Middle panel:</u> von Frey thresholds (g) after vehicle, 2.5 mg/kg morphine, or 2.5 mg/kg morphine + naloxone (1.0 mg/kg, i.p.). <u>Right panel:</u> von Frey thresholds (g) after vehicle, 2.5 mg/kg morphine, or 2.5 mg/kg morphine after 21 days of chronic morphine (2.5 mg/kg/day, i.p.). Wildtype (7 males, 3 females), *Oprm1*-Cre (7 males, 3 females) **(B)** <u>Tail flick test and timeline of experiment for morphine analgesia and tolerance</u>. <u>Left panel:</u> Latency (s) measured after ascending doses of vehicle and morphine (1.25, 2.5, 5, and 10 mg/kg, i.p.). <u>Middle panel:</u> Latency (s) after vehicle, 5 mg/kg morphine, or 5 mg/kg morphine + naloxone (1.0 mg/kg, i.p.). <u>Right panel:</u> Latency (s) after vehicle, 5 mg/kg morphine, or 5 mg/kg morphine after 21 days of chronic morphine (5 mg/kg/day, i.p.). Wildtype (5 males, 5 females), *Oprm1*-Cre (5 males, 4 females). **(C)** <u>Acute lactic acid-induced suppression of operant responding for food pellets</u>. <u>Left panels:</u> Mean±sem pellet intake change score from baseline after injections of lactic acid (0, 0.9, 1.35, and 1.8%, i.p.), morphine (0, 1, 3, and 10 mg/kg, s.c.), and lactic acid (1.8%) plus morphine (0, 1, and 3 mg/kg, s.c). <u>Right panels:</u> Mean±sem percent change from baseline of the data presented on the left panels. Wildtype (4 males, 4 females) and *Oprm1*-Cre (4 males, 3 females) rats.

#### Experiment 3b: Tail flick test

There were no genotype differences in morphine dose response tail flick latencies (Fig. 5B, left). The statistical analysis of latency, which included the between-subjects factor of Genotype and the within-subjects factor of Dose (0, 1.25, 2.5, 5, 10 mg/kg) showed significant effects of Dose (F(4,61.58)=58.1, p<0.001) but no significant effects of Genotype or interaction between the two factors (p>0.1). There were also no genotype differences in response to naloxone or to the analgesic tolerance to morphine (Fig. 5B, middle and right). The statistical analysis of latency, which included the between-subjects factor of Genotype and the within-subjects factor of Condition (Morphine + Naloxone, or Morphine + Tolerance), showed significant effects of Condition for response to naloxone (F(2,23.01)=22.8, p<0.001) and for analgesic tolerance (F(2,33.63)=20.8, p<0.001) but no significant effects of Genotype or interaction between the two factors (p>0.1).

#### Experiment 3c: Lactic acid-induced suppression of operant responding for food

There were no genotype differences in lactic acid concentration response, morphine dose response, and reversal of lactic acid induced suppression of food responding by morphine (Fig. 5C, left panels). Unlike in Experiment 1, in Experiments 4, and 5 there were baseline differences in pellet intake between *Oprm1*-Cre rats and their wildtype littermates (Mean±sem number of pellets per session during the 3 baseline days before lactic acid injections was 106±8 for wildtypes and 133±6 for Oprm1-Cre rats). Thus, we calculated change scores from baseline pellet intake for data presentation and statistical analyses. We also show in Fig. 5C (right panels) the percent change score from baseline.

The statistical analysis of lever press change score for lactic acid, which included the between-subjects factor of Genotype and the within-subjects factor of Dose (0, 0.9, 1.3, 1.8%), showed significant effects of Dose (F(3,39)=17.2, p<0.001). The statistical analysis of lever press change score for morphine, which included the between-subjects factor of Genotype and the within-subjects factor of Morphine Dose (0, 1, 3, 10 mg/kg), showed significant effects of Dose (F(3,39)=58.1, p<0.001). The statistical analysis of pellet intake change score for lactic acid (1.8%) + morphine, which included the between-subjects factor of Genotype and the within-subjects factor of Morphine Dose (0, 1, 3 mg/kg), showed significant effects of F(2,26)=25.0, p>0.001). In all analyses, there were no significant effects of Genotype or interaction between the other factors (p>0.1).

Together, the results of Experiment 3 indicate that the knock-in manipulation had no effect on pain sensitivity to mechanical stimulation, to a noxious thermal stimulus, to acute morphine analgesia, to morphine analgesic tolerance, to suppression of operant responding by lactic acid or morphine, or to reversal of the suppression effect of lactic acid on operant responding by morphine.

### Experiment 4: Effect of Cre-dependent AAV-DIO-Casp3 NAc lesions on acquisition and maintenance of heroin self-administration

#### Food self-administration

##### Acquisition

Bilateral AAV-DIO-Casp3 lesions had no effect on acquisition of food self-administration in *Oprm1*-Cre male and females rats or their wildtype littermates (Fig. 6A, left panels). The statistical analysis of number of pellets, which included the between-subjects factors of Genotype and Sex and the within-subjects factor of Session (1 to 12), showed significant effects of Session (F(11,561)=24.2, p<0.001), but no other significant main or interaction effects (p>0.1).

**Figure 6.**
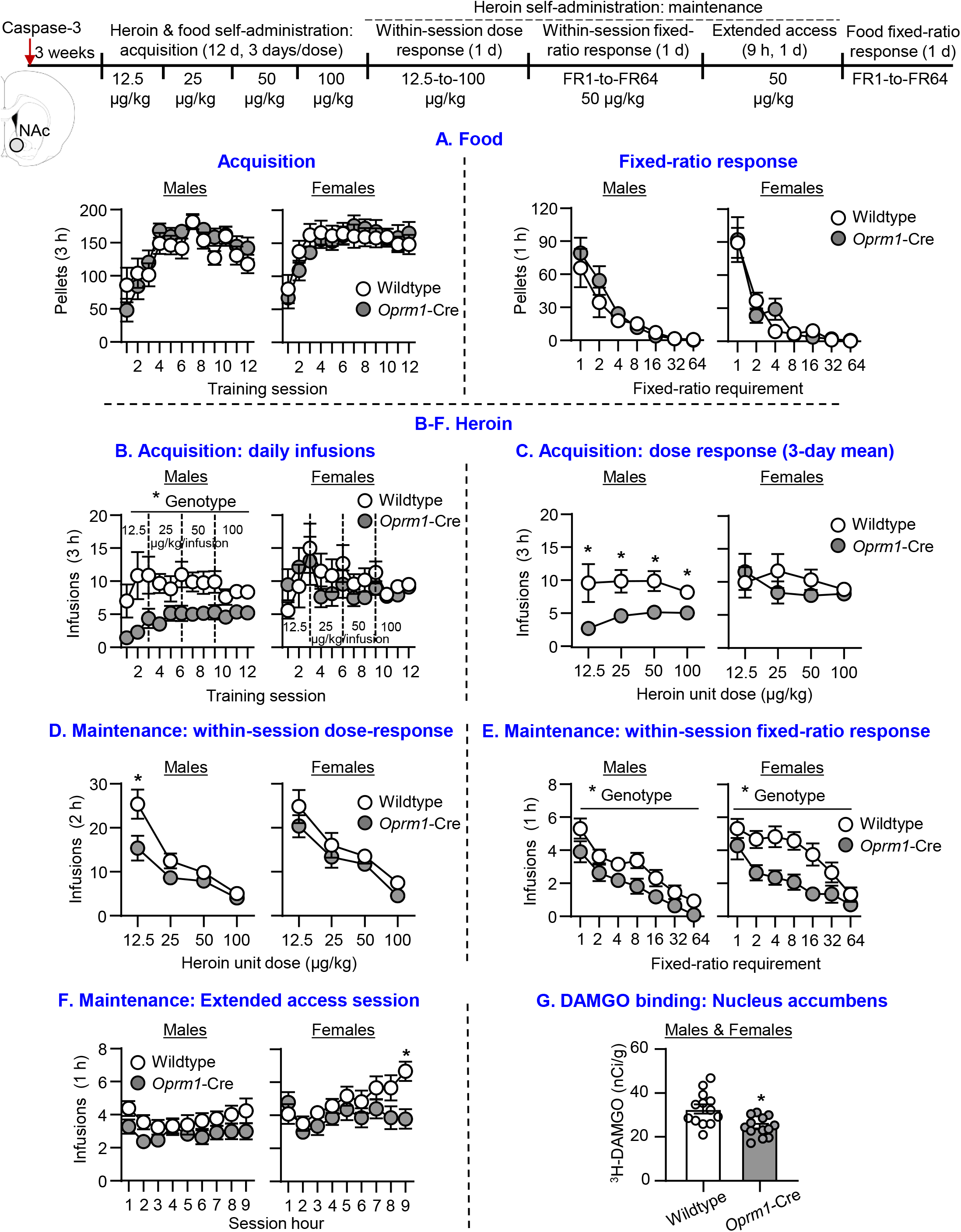
Effect of AAV-DIO-Casp3 NAc lesions on acquisition and maintenance of food and heroin self-administration in heterozygote Oprm1-Cre rats and wildtype littermates. **Food (A)** Left: <u>Acquisition:</u> Mean±sem number of pellets consumed during the 3-h sessions. Wildtype (14 males, 12 females), *Oprm1*-Cre (14 males, 15 females). Right: <u>Within-session fixed-ratio response:</u> Mean±sem number of pellets consumed during the 1-h sessions. Wildtype (8 males, 5 females), *Oprm1*-Cre (8 males, 6 females). **Heroin (B-F): (B)** <u>Acquisition: daily infusions:</u> Mean±sem number of heroin infusions during the 3-h heroin self-administration sessions. **(C)** <u>Acquisition: dose response (3-day mean):</u> Mean±sem of heroin infusions at each unit dose. Wildtype (13 males, 12 females), *Oprm1*-Cre (13 males, 15 females) for **B-C. (D)** <u>Maintenance: within-session dose response:</u> Mean±sem number of heroin infusions at each unit dose. **(E)** <u>Maintenance: within-session fixed-ratio response:</u> Mean±sem number of heroin infusions at each hour of heroin self-administration for each fixed-ratio requirement. **(F)** <u>Maintenance: extended access:</u> Mean±sem number of heroin infusions during the 9-h heroin self-administration session. Wildtype (13 males, 12 females), *Oprm1*-Cre (11 males, 14 females) for **D-F. (G)** [^3^H]DAMGO binding in NAc in wildtype (8 males, 5 females) and *Oprm1*-Cre (8 males, 5 females) rats. Values are calibrated and expressed as nCi/g. * Different from the control hemisphere, p<0.05. * Different from the *Oprm1*-Cre group, p<0.05.

##### Fixed-ratio response

AAV-DIO-Casp3 had no effect on food self-administration under the different FR requirements in *Oprm1*-Cre male and females rats or their wildtype littermates (Fig. 6A, right panels). The statistical analysis of number of pellets, which included the between-subjects factors of Genotype and Sex and the within-subjects factor of FR requirement (1 to 64), showed significant effects of FR requirement (F(6,142)=49.8, p<0.001), but no other significant main or interaction effects (p>0.1).

#### Heroin self-administration

##### Acquisition

AAV-DIO-Casp3 lesions decreased acquisition of heroin self-administration in *Oprm1*-Cre male, but not female rats (Fig. 6B). The statistical analysis of number of daily infusions, which included the between-subjects factors of Genotype and Sex and the within-subjects factor of Heroin Dose (12.5, 25, 50, 100 µg/kg) showed significant effects of Genotype (F(1,49)=5.1, p=0.029) and Sex (F(1,49)=4.1, p=0.049) but no other significant main or interaction effects (p>0.1). For males, a follow-up mixed ANOVA, which included the between-subjects factor of Genotype and the within-subjects factor of Heroin Dose, showed significant effects of Genotype (F(1,24)=8.0, p=0.009) but no significant main effect of Dose or interaction (p>0.1). The same analysis for females did not show any significant effects.

Finally, a mixed ANOVA analysis of the 3-day mean infusions within each Heroin Dose (Fig. 6C), which included the between-subjects factors of Genotype and Sex, and the within-subjects factor of Heroin Dose, showed significant effects of Genotype (F(1,49)=5.1, p=0.029) and Sex (F(1,49)=4.1, p=0.049) but no significant effects of Genotype or interactions between the different factors (p>0.1). Follow up ANOVA within each sex showed a significant effect of Genotype for males (F(1,24)=8.0, p=0.009) but not females (p>0.1).

##### Maintenance: Within-session dose response

AAV-DIO-Casp3 lesions decreased heroin self-administration in *Oprm1*-Cre male but not female rats; this effect was stronger at the lower heroin unit doses (Fig. 6D). The statistical analysis of number of infusions, which included the between-subjects factors of Genotype and Sex, and the within-subjects factor of Heroin Dose (12.5, 25, 50, 100 µg/kg), showed significant effects of Genotype (F(1,46)=4.7, p=0.035), Heroin Dose (F(3,138)=88.2, p<0.001), and Genotype x Heroin dose (F(3,138)=3.1, p=0.029) but no other significant main or interaction effects (p>0.1). For males, a follow-up mixed ANOVA, which included the between-subjects factor of Genotype and the within-subjects factor of Heroin Dose, showed significant effects of Heroin Dose (F(3,66)=49.4, p<0.001) and Genotype x Heroin Dose (F(3,66)=4.6, p=0.006) but no other significant main or interaction effects (p>0.1). The same analysis for females showed a significant effect of Heroin Dose (F(3,75)=40.0, p<0.001), but no significant effects of Genotype or interactions between the two factors (p>0.1).

##### Maintenance: Fixed ratio response

AAV-DIO-Casp3 lesions decreased heroin self-administration in both Oprm1-Cre male and female rats; the lesion effect was stronger at the intermediate FR requirements and appeared stronger in females (Fig. 6E). The statistical analysis of number of infusions, which included the between-subjects factors of Genotype and Sex, and the within-subjects factor of FR requirement (FR1 to FR64), showed significant effects of Genotype (F(1,46)=14.0, p>0.001) and FR requirement (F(6,276)=54.6, p<0.001) but no other significant main or interaction effects (p>0.1). For males, a follow-up mixed ANOVA, which included the between-subjects factor of Genotype and the within-subjects factor of FR requirement, showed significant effects of Genotype (F(1,22)=5.2, p=0.033) and FR requirements (F(6,132)=40.6, p>0.001) but no significant interaction (p>0.1). The same analysis for females showed a significant effect of Genotype (F(1,24)=9.1, p=0.006) and FR requirements (F(6,144)=20.5, p>0.001) but no significant interaction (p>0.1).

##### Maintenance: extended access session (9 h)

AAV-DIO-Casp3 lesions decreased extended access heroin self-administration in *Oprm1-*Cre female, but not male rats (Fig. 6F). The statistical analysis of number of infusions, which included the between-subjects factors of Genotype and Sex and the within-subjects factor of Hour (1 to 9) showed significant effects of Genotype (F(1,46)=5.4, p=0.025), Sex (F(1,46)=8.0, p=0.007), Hour (F(8,365)=5.1, p<0.001), and Genotype x Hour (F(8,365)=2.0, p=0.041) but no other significant main or interaction effects (p>0.1). For males, a follow-up mixed ANOVA, which included the between-subjects factor of Genotype and the within-subjects factor of Hour, showed no significant effects (p>0.05). The same analysis for females showed a significant effect of Hour (F(8,191)=4.4, p>0.001) and Genotype x Hour (F(8,191)=2.6, p=0.010).

##### [^3^H]DAMGO autoradiography

AAV-DIO-Casp3 lesions decreased [^3^H]DAMGO binding in *Oprm1-*Cre male and female rats, but not wildtype littermates (8 males, 5 females/genotype, sex not analyzed independently) (Fig. 6G). The statistical analysis of binding, which included the between-subjects factor of Genotype, showed a significant effect (F(1,24)=11.0, p=0.003).

Together, the results of Experiment 4 indicate that AAV-DIO-Casp3 NAc lesions in *Oprm1*-Cre rats but not wildtype littermates (1) decreased acquisition of heroin self-administration in males but not females, (2) decreased heroin self-administration under an FR1 reinforcement schedule at low but not higher heroin unit doses in males but not females, (3) decreased heroin self-administration when the response requirement were increased in both males and females, (4) decreased heroin self-administration in females but not males when the daily session was increased to 9 h (extended access condition), and (5) decreased NAc [^3^H]DAMGO binding in *Oprm1-Cre* male and female rats. In contrast, NAc Caspase-3 lesions had no effect on acquisition and maintenance of food self-administration under different FR reinforcement schedules in *Oprm1-*Cre rats.

### Experiment 5: Effect of control virus AAV1-DIO-EYFP into NAc on acquisition and maintenance of heroin self-administration

#### Food self-administration

##### Acquisition

AAV-DIO-EYFP had no effect on acquisition of food self-administration in *Oprm1*-Cre rats or their wildtype littermates (Fig. 7A, left panels). The statistical analysis of number of pellets, which included the between-subjects factors of Genotype and Sex and the within-subjects factor of Session (1 to 12), showed significant effects of Session (F(11,286)=6.8, p<0.001), but no significant effects of Genotype or interaction (p>0.1).

**Figure 7.**
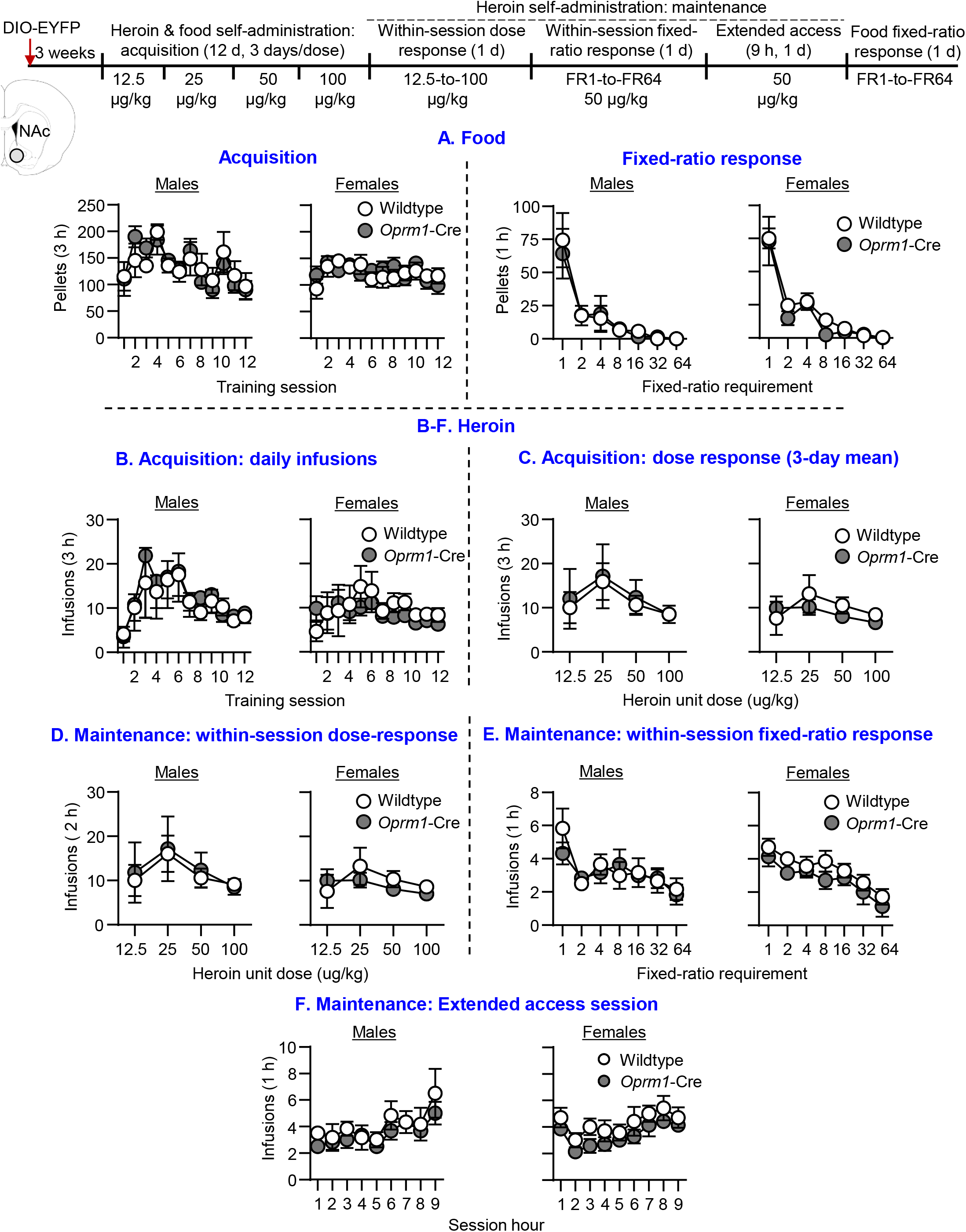
Effect of NAc injections of AAV-DIO-EYFP on acquisition and maintenance of food and heroin self-administration in heterozygote Oprm1-Cre rats and wildtype littermates. **Food (A)** Left: <u>Acquisition:</u> Mean±sem number of pellets consumed during the 3-h sessions. Wildtype (7 males, 8 females), *Oprm1*-Cre (6 males, 7 females). Right: <u>Within-session fixed-ratio response:</u> Mean±sem number of pellets consumed during the 1-h sessions. Wildtype (7 males, 8 females), *Oprm1*-Cre (6 males, 7 females). **Heroin (B-F). (B)** <u>Acquisition: daily infusions:</u> Mean±sem number of heroin infusions during the 3-h heroin self-administration sessions. **(C)** <u>Acquisition: dose response (3-day mean):</u> Mean±sem of heroin infusions for each unit dose. Wildtype (7 males, 8 females), *Oprm1-Cre* (6 males, 7 females) for **(B-C). (D)** <u>Maintenance: within-session dose response:</u> Mean±sem number of heroin infusions for each unit dose. **(E)** <u>Maintenance: within-session fixed-ratio response:</u> Mean±sem number of heroin infusions during each hour of heroin self-administration for each fixed-ratio requirement. **(F)** <u>Maintenance: extended access:</u> Mean±sem number of heroin infusions during the 9-h heroin self-administration session. Wildtype (6 males, 7 females), *Oprm1*-Cre (6 males, 7 females) for **(D-F)**.

##### Fixed ratio response

AAV-DIO-EYFP had no effect on food self-administration under different FR requirements in *Oprm1*-Cre rats or their wildtype littermates (Fig. 7A, right panels). The statistical analysis of number of pellets, which included the between-subjects factor of Genotype and the within-subjects factor of FR requirement (1 to 64), showed significant effects of FR requirement (F(6,156)=50.7, p<0.001), but no significant effects of Genotype or interaction (p>0.1).

#### Heroin self-administration

##### Acquisition

AAV-DIO-EYFP had no effect on acquisition of heroin self-administration in *Oprm1*-Cre rats or their wildtype littermates (Fig. 7B). The statistical analysis of number of daily infusions, which included the between-subjects factor of Genotype and the within-subjects factors of Session (1 to 3) and Heroin Dose (12.5, 25, 50, 100 µg/kg), showed a significant effect of Heroin Dose (F(3,78)=4.7, p=0.005), Session (F(2,52)=4.6, p=0.014), and Session x Heroin Dose (F(6,156)=2.5, p=0.023) but no significant effects of Genotype or interactions between Genotype and the other factors (p>0.1). A mixed ANOVA analysis of the 3-day mean infusions within each Heroin Dose (Fig. 7C), which included the between-subjects factor of Genotype and the within-subjects factors of Heroin Dose, showed a significant effect of Heroin Dose (F(3,78)=7.0, p<0.001) but no significant effects of Genotype or interaction (p>0.1).

##### Maintenance: Within-session dose response

AAV-DIO-EYFP had no effect on heroin self-administration in Oprm1-Cre rats or wildtype littermates (Fig. 7D). The statistical analysis of number of infusions, which included the between-subjects factor of Genotype and the within-subjects factor of Heroin Dose (12.5, 25, 50, 100 µg/kg), showed significant effects of Heroin Dose (F(3,72)=49.1, p<0.001) but no significant effect of Genotype or interaction (p>0.1).

##### Maintenance: Fixed ratio response

AAV-DIO-EYFP had no effect on heroin self-administration in *Oprm1-*Cre rats or wildtype littermates (Fig. 7E). The statistical analysis of number of infusions, which included the between-subjects factor of Genotype and the within-subjects factor of FR requirement (FR1 to FR64), showed significant effects of FR requirement (F(6,144)=16.4, p<0.001) but no significant effect of Genotype or interaction (p>0.1).

##### Maintenance: extended access session (9 h)

AAV-DIO-EYFP had no effect on extended access heroin self-administration in *Oprm1-*Cre rats or their wildtype littermates (Fig. 7F). The statistical analysis of number of infusions, which included the between-subjects factor of Genotype and Sex and the within-subjects factor of Hour (1 to 9), showed significant effects of Hour (F(8,192)=10.6, p<0.001) but no significant effects of Genotype or interaction between Genotype and the other factors(p>0.1).

Together, the results of Experiment 5 indicate that NAc DIO-EYFP had no effect on any of the heroin self-administration behaviors in either *Oprm1-*Cre rats or their wildtype littermates, suggesting that non-lesioning Cre-dependent manipulations do not affect MOR expression or opioid-mediated behaviors.

## Discussion

We performed anatomical and behavioral characterization of a novel CRISPR-mediated knock-in rat that co-expresses Cre-recombinase and MOR under the endogenous *Oprm1* gene promoter. We report four main findings. First, the knock-in manipulation had no effect on *Oprm1* mRNA expression in dHipp, NAc, and DS, MOR function in NAc and DS, and MOR density in NAc; however, MOR density in DS was higher in *Oprm1*-Cre rats. Second, insertion of T2A-iCre resulted in >95% colocalization of *Oprm1* and Cre in *Oprm1*-Cre rats in NAc, DS, and dHipp; additionally, Cre was not detected in wildtype littermates. Third, the knock-in of Cre into the *Oprm1* gene had no effect on operant responding for food pellets or MOR-mediated behaviors, including pain sensitivity, morphine analgesia and tolerance, heroin self-administration, and heroin relapse-related behaviors. Fourth, lesions of NAc MOR-expressing cells using a Cre-dependent AAV-DIO-Casp3 virus decreased *Oprm1* mRNA expression and MOR density in *Oprm1*-Cre rats but not wildtype littermates. Additionally, the lesion had sex-specific effects on acquisition and maintenance of heroin self-administration. Together, these results indicate that MOR expression and function are preserved in the novel *Oprm1*-Cre knock-in rat, and that we can selectively target and manipulate brain MOR-expressing cells to study MOR-mediated behaviors.

### The *Oprm1* knock-in manipulation had no effect on several behavioral measures and MOR expression and function

We behaviorally characterized the *Oprm1* knock-in rats in three ways: (1) operant responding for food pellets, (2) pain sensitivity and morphine analgesia using three different methods to evaluate pain-related behaviors, and (3) heroin self-administration and relapse-related behaviors.

*Oprm1-*Cre rats did not differ from their wildtype littermates in acquisition of food self-administration and sensitivity to increasing the effort (response requirement) to obtain the food reinforcer. *Oprm1-*Cre rats also did not differ from their wildtype littermates in three different pain-related assays: tail flick test, von Frey test, and lactic acid inhibition of operant responding for food. *Oprm1-*Cre rats also did not differ from their wildtype littermates in sensitivity to morphine-induced analgesia, the development of morphine tolerance, and reversal of morphine’s analgesic effects by naloxone. Finally, *Oprm1-*Cre rats also did not differ from their wildtype littermates in acquisition and maintenance of heroin self-administration, and lever responding in three commonly used relapse-related measures (Shalev et al., 2002; Reiner et al., 2019): extinction responding, context-induced reinstatement, and reacquisition after extinction. Together, our initial behavioral characterization indicates that the *Oprm1* knock-in manipulation did not alter normal operant learning, sensitivity to food reward, and prototypical MOR-mediated behaviors like opioid analgesia, opioid tolerance, and opioid self-administration.

### Methodological considerations

The insertion of 2A-Cre into the coding sequence of Oprm1 may negatively affect how MOR is expressed at both the RNA and protein level leading to altered MOR functionality. Our anatomical (and behavioral) characterization suggests that the CRISPR-based insertion of T2A-iCre did not change *Oprm1* expression or function. However, this conclusion should be interpreted with caution because of several technical issues. Specifically, in our HCR FISH experiment using an *Oprm1*-specific probe on heterozygous *Oprm1*-Cre rats, we did not detect statistically significant differences between *Oprm1*-Cre rats and wildtype littermates in number of *Oprm1*-postive cells in dHipp, NAc, or DS. However, our experiment did not address whether the heterozygous genotype results in transcriptional changes at the cellular level. Additionally, because we cannot differentiate the transgenic mRNA from the endogenous mRNA that is normally expressed in wildtype rats, we are unable to conclude that the transgene expression is restricted to cells that would otherwise express endogenous *Oprm1*. Because of this, we cannot rule out the possibility that the Cre insertion will affect *Oprm1* expression in terms of spatial distribution, temporal dynamics, and response to external stimuli like opioid drugs and stress. Additionally, the reasons for the somewhat higher [^3^H]DAMGO binding but not activity in DS are unknown. Finally, while we report no differences in HCR FISH or [^35^S]GTPγS activity assays, and pharmacology/behavioral assays, it remains to be determined if these “non-phenotypes” generalize to homozygote *Oprm1*-Cre rats.

### Effect of lesions of MOR-expressing cells in NAc on heroin self-administration in males and females

Studies using systemic injections of selective or preferential (e.g., naloxone, naltrexone) antagonists of MOR and other opioid receptors indicate that activation of MOR is critical for the reinforcing effects of opioid agonists in the drug self-administration procedure in male rodents and monkeys (Goldberg et al., 1971; Ettenberg et al., 1982; Wise, 1989; Mello and Negus, 1996). There is also evidence from studies using systemic injections of opioid receptor antagonists in male rats that activation of MOR is critical for reinstatement of opioid seeking induced by drug priming, and drug cues and contexts (Shaham and Stewart, 1996; Bossert et al., 2019; Reiner et al., 2019).

Studies using site-specific injections of lipophobic preferential MOR antagonists (methyl naltrexone or methyl naloxonium) into different brain areas indicate that the critical site of action for the systemic effect of MOR antagonists on opioid self-administration in male rats is NAc; there is also evidence for a role of ventral tegmental area and periaqueductal gray (Vaccarino and Corrigall, 1987; Corrigall and Vaccarino, 1988). In contrast, the site of action for the effect of systemic preferential MOR antagonists on reinstatement of opioid seeking is unknown (Reiner et al., 2019), and the role of brain MOR in opioid self-administration and reinstatement in female rats is unknown.

Based on the literature described above, we used the new *Oprm1*-Cre rat to determine the role of MOR-expressing cells in NAc in initiation and maintenance of heroin self-administration, using a within-subjects ascending heroin dose-response curve procedure (Stewart et al., 1996). We found that injections of Cre-dependent AAV1-EF1a-Flex-taCasp3-TEVP into NAc of *Oprm1-*Cre rats, which selectively lesion MOR-expressing cells, had sex-specific effects on heroin self-administration. During the acquisition phase, the lesions selectively decreased heroin self-administration at all four heroin doses in males but not females. During the maintenance phase, the lesions decreased responding for the lowest heroin dose in males but not females, decreased extended-access heroin self-administration in females but not males, and had a more pronounced inhibitory effect on the effort (response requirement) to self-administer heroin in males than in females.

Our results suggest mechanistic differences in the role of MOR-expressing cells in heroin seeking and taking. This notion is supported by results from our recent studies on the sex-dependent effects of TRV130 (a selective partial agonist of MOR) and BU08028 (a mixed partial agonist of MOR and nociceptin/orphanin/FQ peptide receptor) on context-induced reinstatement and reacquisition of heroin and oxycodone self-administration (Bossert et al., 2020; Bossert et al., 2022). These sex-dependent effects in opioid relapse occurred even though we did not observe sex differences in opioid self-administration in our previous studies and in the current study. A question for future research is the mechanism/s of the potential sex-specific role of NAc MOR-expressing cells in heroin self-administration. One possibility is sex-specific modulation of striatal MOR by local estrogen and progesterone receptors (Mansour et al., 1988; Tobiansky et al., 2018), which play a role in striatal-dependent (Kalivas and Stewart, 1991) locomotor sensitization to opioid drugs (Craft et al., 2006).

Overall, our results with the new *Oprm1*-Cre rats extend results from previous pharmacological studies on the role of NAc MOR in heroin self-administration in male rats and suggest that MOR-expressing cells in this region play distinct roles in heroin reinforcement in males and females. However, from a statistical perspective, an interpretation caveat of our data is that our conclusions are based on post-hoc analysis within each sex, without a statistically significant sex by genotype interaction in the initial full factorial ANOVAs.

### Conclusions

We anatomically and behaviorally validated a CRISPR-based *Oprm1*-Cre knock-in rat that allows cell-type specific genetic access to measure and manipulate brain MOR-expressing cells. We used the *Oprm1*-Cre rats to show a potential sex-specific role of NAc MOR-expressing cells in heroin self-administration. The ability to specifically alter the neurons expressing MOR, and to target MOR itself and its downstream signaling, offers significant mechanistic advantages given the lack of receptor selectivity of many of the endogenous opioid ligands and of several classical opioid agonist and antagonist drugs (Heyman et al., 1986; Mansour et al., 1995b). The new *Oprm1*-Cre rats can be used to study both the general role and the sex-specific role of brain MOR-expressing cells in rat models of opioid addiction and pain-related behaviors, and other opioid-mediated behaviors.

## Acknowledgments

The research was supported by funds to the Intramural Research Program of NIDA (ZIA-DA000434-22 and ZIA-DA000069), and NIDA U01DA043098, ONR 00014-19-1-2149, and HDRF (to H. Akil and S.J. Watson).

## References

Adhikary S, Caprioli D, Venniro M, Kallenberger P, Shaham Y, Bossert JM (2017) Incubation of extinction responding and cue-induced reinstatement, but not context-or drug priming-induced reinstatement, after withdrawal from methamphetamine. Addict Biol 22:977–990.

Akil H, Watson SJ, Young E, Lewis ME, Khachaturian H, Walker JM (1984) Endogenous opioids: biology and function. Annu Rev Neurosci 7:223–255.

Anderson KR et al. (2018) CRISPR off-target analysis in genetically engineered rats and mice. Nat Methods 15:512–514.

Bailly J, Del Rossi N, Runtz L, Li JJ, Park D, Scherrer G, Tanti A, Birling MC, Darcq E, Kieffer BL (2020) Targeting Morphine-Responsive Neurons: Generation of a Knock-In Mouse Line Expressing Cre Recombinase from the Mu-Opioid Receptor Gene Locus. eNeuro 7.

Baldwin AN, Banks ML, Marsh SA, Townsend EA, Venniro M, Shaham Y, Negus SS (2022) Acute pain-related depression of operant responding maintained by social interaction or food in male and female rats. Psychopharmacology (Berl).

Basila M, Kelley ML, Smith AVB (2017) Minimal 2’-O-methyl phosphorothioate linkage modification pattern of synthetic guide RNAs for increased stability and efficient CRISPR-Cas9 gene editing avoiding cellular toxicity. PLoS One 12:1–19.

Bossert J, Liu S, Lu L, Shaham Y (2004) A role of ventral tegmental area glutamate in contextual cue-induced relapse to heroin seeking. J Neurosci 24:10726–10730.

Bossert JM, Marchant NJ, Calu DJ, Shaham Y (2013) The reinstatement model of drug relapse: recent neurobiological findings, emerging research topics, and translational research. Psychopharmacology 229:453–476.

Bossert JM, Adhikary S, St Laurent R, Marchant NJ, Wang HL, Morales M, Shaham Y (2016) Role of projections from ventral subiculum to nucleus accumbens shell in context-induced reinstatement of heroin seeking in rats. Psychopharmacology 233:1991–2004.

Bossert JM, Hoots JK, Fredriksson I, Adhikary S, Zhang M, Venniro M, Shaham Y (2019) Role of mu, but not delta or kappa, opioid receptors in context-induced reinstatement of oxycodone seeking. Eur J Neurosci 50:2075–2085.

Bossert JM, Townsend EA, Altidor LK, Fredriksson I, Shekara A, Husbands S, Sulima A, Rice KC, Banks ML, Shaham Y (2022) Sex differences in the effect of chronic delivery of the buprenorphine analogue BU08028 on heroin relapse and choice in a rat model of opioid maintenance. Br J Pharmacol 179:227–241.

Bossert JM, Kiyatkin E, Korah H, Hoots JK, Afzal A, Perekopskiy D, Thomas S, Fredriksson I, Blough BE, Negus SS, Epstein DH, Shaham Y (2020) In a rat model of opioid maintenance, the G-protein-biased MOR agonist TRV130 decreases relapse to oxycodone seeking and taking, and prevents oxycodone-induced brain hypoxia. Biol Psychiatry 88:935–944.

Brinkman EK, Chen T, Amendola M, van Steensel B (2014) Easy quantitative assessment of genome editing by sequence trace decomposition. Nucleic Acids Res 42:1–8.

Caprioli D, Venniro M, Zeric T, Li X, Adhikary S, Madangopal R, Marchant NJ, Lucantonio F, Schoenbaum G, Bossert JM, Shaham Y (2015) Effect of the novel positive allosteric modulator of metabotropic glutamate receptor 2 AZD8529 on incubation of methamphetamine craving after prolonged voluntary abstinence in a rat model. Biol Psychiatry 78:463–473.

Charbogne P et al. (2017) Mu Opioid Receptors in Gamma-Aminobutyric Acidergic Forebrain Neurons Moderate Motivation for Heroin and Palatable Food. Biol Psychiatry 81:778–788.

Choi HMT, Schwarzkopf M, Fornace ME, Acharya A, Artavanis G, Stegmaier J, Cunha A, Pierce NA (2018) Third-generation in situ hybridization chain reaction: multiplexed, quantitative, sensitive, versatile, robust. Development 145:1–10.

Chow JJ, Beacher N, Chabot JM, Oke M, Venniro M, Lin DT, Shaham Y (2022) Characterization of operant social interaction in rats: effects of access duration, effort, peer familiarity, housing conditions, and choice between social interaction versus food or remifentanil. Psychopharmacology (in press).

Concordet JP, Haeussler M (2018) CRISPOR: intuitive guide selection for CRISPR/Cas9 genome editing experiments and screens. Nucleic Acids Res 46:W242–W245.

Corrigall WA, Vaccarino FJ (1988) Antagonist treatment in nucleus accumbens or periaqueductal grey affects heroin self-administration. Pharmacol Biochem Behav 30:443–450.

Craft RM, Clark JL, Hart SP, Pinckney MK (2006) Sex differences in locomotor effects of morphine in the rat. Pharmacol Biochem Behav 85:850–858.

Darcq E, Kieffer BL (2018) Opioid receptors: drivers to addiction? Nat Rev Neurosci 19:499–514.

Deroche V, Le Moal M, Piazza PV (1999) Cocaine self-administration increases the incentive motivational properties of the drug in rats. Eur J Neurosci 11:2731–2736.

Doench JG, Fusi N, Sullender M, Hegde M, Vaimberg EW, Donovan KF, Smith I, Tothova Z, Wilen C, Orchard R, Virgin HW, Listgarten J, Root DE (2016) Optimized sgRNA design to maximize activity and minimize off-target effects of CRISPR-Cas9. Nat Biotechnol 34:184–191.

Emery MA, Akil H (2020) Endogenous Opioids at the Intersection of Opioid Addiction, Pain, and Depression: The Search for a Precision Medicine Approach. Annu Rev Neurosci 43:355–374.

Ettenberg A, Pettit HO, Bloom FE, Koob GF (1982) Heroin and cocaine intravenous self-administration in rats: mediation by separate neural systems. Psychopharmacology (Berl) 78:204–209.

Filipiak WE, Saunders TL (2006) Advances in transgenic rat production. Transgenic Res 15:673–686.

Filipiak WE, Hughes ED, Gavrilina GB, LaForest AK, Saunders TL (2019) Next Generation Transgenic Rat Model Production. Methods Mol Biol 2018:97–114.

Fredriksson I, Applebey SV, Minier-Toribio A, Shekara A, Bossert JM, Shaham Y (2020) Effect of the dopamine stabilizer (-)-OSU6162 on potentiated incubation of opioid craving after electric barrier-induced voluntary abstinence. Neuropsychopharmacology 45:770–779.

Fredriksson I, Adhikary S, Steensland P, Vendruscolo LF, Bonci A, Shaham Y, Bossert JM (2017) Prior Exposure to Alcohol Has No Effect on Cocaine Self-Administration and Relapse in Rats: Evidence from a Rat Model that Does Not Support the Gateway Hypothesis. Neuropsychopharmacology 42:1001–1011.

Gardon O, Faget L, Chu Sin Chung P, Matifas A, Massotte D, Kieffer BL (2014) Expression of mu opioid receptor in dorsal diencephalic conduction system: new insights for the medial habenula. Neuroscience 277:595–609.

Gelman A, Hill J (2006) Data analysis using regression and multilevel/hierarchical models: Cambridge University Press.

Goldberg SR, Woods JH, Schuster CR (1971) Nalorphine-induced changes in morphine self-administration in rhesus monkeys. J Pharmacol Exp Ther 176:464–471.

Heyman JS, Koslo RJ, Mosberg HI, Tallarida RJ, Porreca F (1986) Estimation of the affinity of naloxone at supraspinal and spinal opioid receptors in vivo: studies with receptor selective agonists. Life Sci 39:1795–1803.

Highfield D, Mead A, Grimm J, Rocha B, Shaham Y (2002) Reinstatement of cocaine seeking in 129X1/SvJ mice: effects of cocaine priming, cocaine cues and food deprivation. Psychopharmacology 161:417–424.

Jaffe JH (1990) Drug addiction and drug abuse. In: Goodman & Gilman’s the pharmacological basis of therapeutics, 8th Edition (Gilman AG, Rall TW, Nies AS, Taylor P, eds), pp 522–573. New York: Pergamon Press.

Kalivas PW, Stewart J (1991) Dopamine transmission in the initiation and expression of drug- and stress-induced sensitization of motor activity. Brain Res Rev 16:223–244.

Khoo SY, Gibson GD, Prasad AA, McNally GP (2017) How contexts promote and prevent relapse to drug seeking. Genes Brain Behav 16:185–204.

Kim JH, Lee SR, Li LH, Park HJ, Park JH, Lee KY, Kim MK, Shin BA, Choi SY (2011) High cleavage efficiency of a 2A peptide derived from porcine teschovirus-1 in human cell lines, zebrafish and mice. PLoS One 6:1–8.

Koob GF (1992) Neural mechanisms of drug reinforcement. Ann N Y Acad Sci 654:171–191.

Kumar V, Krolewski DM, Hebda-Bauer EK, Parsegian A, Martin B, Foltz M, Akil H, Watson SJ (2021) Optimization and evaluation of fluorescence in situ hybridization chain reaction in cleared fresh-frozen brain tissues. Brain Struct Funct 226:481–499.

Mansour A, Fox CA, Akil H, Watson SJ (1995a) Opioid-receptor mRNA expression in the rat CNS: anatomical and functional implications. Trends Neurosci 18:22–29.

Mansour A, Khachaturian H, Lewis ME, Akil H, Watson SJ (1988) Anatomy of CNS opioid receptors. Trends Neurosci 11:308–314.

Mansour A, Hoversten MT, Taylor LP, Watson SJ, Akil H (1995b) The cloned mu, delta and kappa receptors and their endogenous ligands: evidence for two opioid peptide recognition cores. Brain Res 700:89–98.

Marchant NJ, Campbell EJ, Pelloux Y, Bossert JM, Shaham Y (2019) Context-induced relapse after extinction versus punishment: similarities and differences. Psychopharmacology 236:439–448.

Marchant NJ, Campbell EJ, Whitaker LR, Harvey BK, Kaganovsky K, Adhikary S, Hope BT, Heins RC, Prisinzano TE, Vardy E, Bonci A, Bossert JM, Shaham Y (2016) Role of ventral subiculum in context-induced relapse to alcohol seeking after punishment-imposed abstinence. J Neurosci 36:3281–3294.

McBurney MW, Fournier S, Jardine K, Sutherland L (1994) Intragenic regions of the murine Pgk-1 locus enhance integration of transfected DNAs into genomes of embryonal carcinoma cells. Somat Cell Mol Genet 20:515–528.

Mello NK, Negus SS (1996) Preclinical evaluation of pharmacotherapies for treatment of cocaine and opioid abuse using drug self-administration procedures. Neuropsychopharmacology 14:375–424.

Paxinos G, Watson C (2007) The rat brain in stereotaxic coordinates. Sixth edition., 6 Edition. San Diego, CA: Academic Press.

Quadros RM et al. (2017) Easi-CRISPR: a robust method for one-step generation of mice carrying conditional and insertion alleles using long ssDNA donors and CRISPR ribonucleoproteins. Genome Biol 18:1–15.

Reiner DJ, Fredriksson I, Lofaro OM, Bossert JM, Shaham Y (2019) Relapse to opioid seeking in rat models: behavior, pharmacology and circuits. Neuropsychopharmacology 44:465–477.

Reiner DJ, Lofaro OM, Applebey SV, Korah H, Venniro M, Cifani C, Bossert JM, Shaham Y (2020) Role of projections between piriform cortex and orbitofrontal cortex in relapse to fentanyl seeking after palatable food choice-induced voluntary abstinence. J Neurosci 40:2485–2497.

Reiner DJ, Townsend EA, Orihuel J, Applebey SV, Claypool SM, Banks ML, Shaham Y, Negus SS (2021) Lack of effect of different pain-related manipulations on opioid self-administration, reinstatement of opioid seeking, and opioid choice in rats. Psychopharmacology (Berl) 238:1885–1897.

Schindelin J, Arganda-Carreras I, Frise E, Kaynig V, Longair M, Pietzsch T, Preibisch S, Rueden C, Saalfeld S, Schmid B, Tinevez JY, White DJ, Hartenstein V, Eliceiri K, Tomancak P, Cardona A (2012) Fiji: an open-source platform for biological-image analysis. Nat Methods 9:676–682.

Schneider CA, Rasband WS, Eliceiri KW (2012) NIH Image to ImageJ: 25 years of image analysis. Nat Methods 9:671–675.

Shaham Y, Stewart J (1996) Effects of opioid and dopamine receptor antagonists on relapse induced by stress and re-exposure to heroin in rats. Psychopharmacology 125:385–391.

Shaham Y, Shalev U, Lu L, De Wit H, Stewart J (2003) The reinstatement model of drug relapse: history, methodology and major findings. Psychopharmacology 168:3–20.

Shalev U, Grimm J, Shaham Y (2002) Neurobiology of relapse to heroin and cocaine seeking: A review. Pharmacol Rev 54:1–42.

Shimshek DR, Kim J, Hubner MR, Spergel DJ, Buchholz F, Casanova E, Stewart AF, Seeburg PH, Sprengel R (2002) Codon-improved Cre recombinase (iCre) expression in the mouse. Genesis 32:19–26.

Slaymaker IM, Gao L, Zetsche B, Scott DA, Yan WX, Zhang F (2016) Rationally engineered Cas9 nucleases with improved specificity. Science 351:84–88.

Stewart J, Woodside B, Shaham Y (1996) Ovarian hormones do not affect the initiation and maintenance of intravenous self-administration of heroin in the female rat. Psychobiology 24:154–159.

Takahashi YK, Batchelor HM, Liu B, Khanna A, Morales M, Schoenbaum G (2017) Dopamine neurons respond to errors in the prediction of sensory features of expected rewards. Neuron 95:1395–1405 e1393.

Terrier J, Luscher C, Pascoli V (2016) Cell-type specific insertion of GluA2-lacking AMPARs with cocaine exposure leading to sensitization, cue-induced seeking, and incubation of craving. Neuropsychopharmacology 41:1779–1789.

Tobiansky DJ, Wallin-Miller KG, Floresco SB, Wood RI, Soma KK (2018) Androgen Regulation of the Mesocorticolimbic System and Executive Function. Front Endocrinol 9:1–18.

Vaccarino FJ, Corrigall WA (1987) Effects of opiate antagonist treatment into either the periaqueductal grey or nucleus accumbens on heroin-induced locomotor activation. Brain Res Bull 19:545–549.

Vaccarino FJ, Bloom FE, Koob GF (1985) Blockade of nucleus accumbens opiate receptors attenuates intravenous heroin reward in the rat. Psychopharmacology (Berl) 86:37–42.

Venniro M, Caprioli D, Shaham Y (2016) Animal models of drug relapse and craving: From drug priming-induced reinstatement to incubation of craving after voluntary abstinence. Prog Brain Res 224:25–52.

Venniro M, Zhang M, Shaham Y, Caprioli D (2017) Incubation of methamphetamine but not heroin craving after voluntary abstinence in male and female rats. Neuropsychopharmacology 42:1126–1135.

Wang S, Lim G, Yang L, Zeng Q, Sung B, Jeevendra Martyn JA, Mao J (2005) A rat model of unilateral hindpaw burn injury: slowly developing rightwards shift of the morphine dose-response curve. Pain 116:87–95.

Weibel R, Reiss D, Karchewski L, Gardon O, Matifas A, Filliol D, Becker JA, Wood JN, Kieffer BL, Gaveriaux-Ruff C (2013) Mu opioid receptors on primary afferent nav1.8 neurons contribute to opiate-induced analgesia: insight from conditional knockout mice. PLoS One 8:e74706.

Wise RA (1989) Opiate reward: sites and substrates. Neurosci Biobehav Rev 13:129–133.

Wolf ME (2016) Synaptic mechanisms underlying persistent cocaine craving. Nat Rev Neurosci 17:351–365.

