## Supplemental Tables 1 & 2 for "Effect of selective lesions of nucleus accumbens μ-opioid receptor-expressing cells on heroin self-administration in male and female rats: a study with novel *Oprm1-Cre* knock-in rats"

10-31-2022

Table of content

Supplementary Table 1. Sequence of donor DNA and split-initiator DNA probe (HCR version 2.0) sequences (5’ to 3’)

Supplementary Table 2. Statistical analyses

**Table S1A**. Sequence of Donor DNA for Generation of *Oprm1*-Cre knock-in rat

| 5' arm of homology (nucleotides 63-98 are *oprm1* exon 4) | CATCAATGTGCTCTCTAATGAGACCCCAGAACTCACTATCTTCACTCTTTCTCTTCTTTCAGCTAGAAAATCTGGAGGCAGAAACTGCTCCATTGCCC |
| --- | --- |
| Gly-Ser-Gly linker and T2A self-cleaving peptide sequence (lower case) | GGTTCTGGCgagggcagaggaagtcttctaacatgcggtgacgtggaggagaatcccggccct |
| iCre recombinase coding sequence | GTGCCCAAGAAGAAGAGGAAAGTCTCCAACCTGCTGACTGTGCACCAAAACCTGCCTGCCCTCCCTGTGGATGCCACCTCTGATGAAGTCAGGAAGAACCTGATGGACATGTTCAGGGACAGGCAGGCCTTCTCTGAACACACCTGGAAGATGCTCCTGTCTGTGTGCAGATCCTGGGCTGCCTGGTGCAAGCTGAACAACAGGAAATGGTTCCCTGCTGAACCTGAGGATGTGAGGGACTACCTCCTGTACCTGCAAGCCAGAGGCCTGGCTGTGAAGACCATCCAACAGCACCTGGGCCAGCTCAACATGCTGCACAGGAGATCTGGCCTGCCTCGCCCTTCTGACTCCAATGCTGTGTCCCTGGTGATGAGGAGAATCAGAAAGGAGAATGTGGATGCTGGGGAGAGAGCCAAGCAGGCCCTGGCCTTTGAACGCACTGACTTTGACCAAGTCAGATCCCTGATGGAGAACTCTGACAGATGCCAGGACATCAGGAACCTGGCCTTCCTGGGCATTGCCTACAACACCCTGCTGCGCATTGCCGAAATTGCCAGAATCAGAGTGAAGGACATCTCCCGCACCGATGGTGGGAGAATGCTGATCCACATTGGCAGGACCAAGACCCTGGTGTCCACAGCTGGTGTGGAGAAGGCCCTGTCCCTGGGGGTTACCAAGCTGGTGGAGAGATGGATCTCTGTGTCTGGTGTGGCTGATGACCCCAACAACTACCTGTTCTGCCGGGTCAGAAAGAATGGTGTGGCTGCCCCTTCTGCCACCTCCCAACTGTCCACCCGGGCCCTGGAAGGGATCTTTGAGGCCACCCACCGCCTGATCTATGGTGCCAAGGATGACTCTGGGCAGAGATACCTGGCCTGGTCTGGCCACTCTGCCAGAGTGGGTGCTGCCAGGGACATGGCCAGGGCTGGTGTGTCCATCCCTGAAATCATGCAGGCTGGTGGCTGGACCAATGTGAACATTGTGATGAACTACATCAGAAACCTGGACTCTGAGACTGGGGCCATGGTGAGGCTGCTCGAGGATGGGGAC |
| Oprm1 termination codon and 3' arm of homology from *oprm1* exon 4 | TAACTGGGTCTCACACCATCCAGACCCTCGCTAAGCTTAGAGGCCGCCATCTACGTGGAATCAGGTTGCTGTCAGGGTGTGTGGGAGGCTCTGGTTTCCTG |

**Table S1B. Oprm1 Probe:** Seq ID - NM_013071.2; HCR amplifier: B2-AlexaFluor647

| **Odd** | **1st half of Initiator I1 + Spacer (AA) + Probe Sequence** | **Even** | **Probe Sequence + Spacer (TA) + 2nd half of Initiator I1** |
| --- | --- | --- | --- |
| 1 | CCTCgTAAATCCTCATCAAAGACACTCTGAAAGGGCAGTGTACTG | 2 | GAAGGGCCATGTTCCCATCAGGTAGAAATCATCCAgTAAACCgCC |
| 3 | CCTCgTAAATCCTCATCAAATCACGATCTTGCAGAGGATGGTTCC | 4 | TGAACATGTTGTAGTAATCTATTGAAAATCATCCAgTAAACCgCC |
| 5 | CCTCgTAAATCCTCATCAAAATGGTGCAGAGGGTGAATATGCTGG | 6 | CAGACAGCAATGTAGCGGTCCACGCAAATCATCCAgTAAACCgCC |
| 7 | CCTCgTAAATCCTCATCAAAAAGAGAGGATCCAGTTGCAGACGTT | 8 | TGAACATTACAGGCAGACCGATGGCAAATCATCCAgTAAACCgCC |
| 9 | CCTCgTAAATCCTCATCAAACCCTGCCTGTATTTTGTGGTTGCCA | 10 | GAGAACGTGAGGGTGCAATCTATGGAAATCATCCAgTAAACCgCC |
| 11 | CCTCgTAAATCCTCATCAAAGTTCTCCCAGTACCAGGTTGGGTGG | 12 | GAAGATAAAGACACAGATTTTGAGCAAATCATCCAgTAAACCgCC |
| 13 | CCTCgTAAATCCTCATCAAATGATGAGGACCGGCATGATGAAAGC | 14 | AGATCATCAGGCCGTAACACACAGTAAATCATCCAgTAAACCgCC |
| 15 | CCTCgTAAATCCTCATCAAAAGCATGCGAACGCTCTTGAGTCGTA | 16 | TTCCTGTCCTTTTCTTTGGAGCCCGAAATCATCCAgTAAACCgCC |
| 17 | CCTCgTAAATCCTCATCAAACACCATCCGGGTGATCCTGCGCAGA | 18 | GACGATAAATACAGCCACGACCACCAAATCATCCAgTAAACCgCC |
| 19 | CCTCgTAAATCCTCATCAAAGAAACGGTCTGAAATGTGGTTTCTG | 20 | TAACCCAAAGCAATGCAGAAGTGCCAAATCATCCAgTAAACCgCC |
| 21 | CCTCgTAAATCCTCATCAAAAACTGGATTCAGGCAGCTGTTCGTG | 22 | GAAGTTTTCATCCAGGAAGGCGTAAAAATCATCCAgTAAACCgCC |
| 23 | CCTCgTAAATCCTCATCAAATGCAGAACTCTCTGAAGCATCGCTT | 24 | GCTGTTCGATCGTGGACGAGGTTGGAAATCATCCAgTAAACCgCC |
| 25 | CCTCgTAAATCCTCATCAAATTCTGACGGACTCGAGTGGAGTTTT | 26 | TTAGCCGTGGAGGGATGTTCCCTAGAAATCATCCAgTAAACCgCC |

**Table S1C. iCre Probe:** Seq ID - AY056050.1; HCR amplifier: B3-AlexaFluor546

| **Odd** | **1st half of Initiator I1 + Spacer (TT) + Probe Sequence** | **Even** | **Probe Sequence + Spacer (TT) + 2nd half of Initiator I1** |
| --- | --- | --- | --- |
| 1 | gTCCCTgCCTCTATATCTTTCTTGGGCACCATGGTGGACAAGCTT | 2 | CAGCAGGTTGGAGACTTTCCTCTTCTTCCACTCAACTTTAACCCg |
| 3 | gTCCCTgCCTCTATATCTTTGGGCAGGCAGGTTTTGGTGCACAGT | 4 | CTTCATCAGAGGTGGCATCCACAGGTTCCACTCAACTTTAACCCg |
| 5 | gTCCCTgCCTCTATATCTTTAACATGTCCATCAGGTTCTTCCTGA | 6 | TGTTCAGAGAAGGCCTGCCTGTCCCTTCCACTCAACTTTAACCCg |
| 7 | gTCCCTgCCTCTATATCTTTCACAGACAGGAGCATCTTCCAGGTG | 8 | CTTGCACCAGGCAGCCCAGGATCTGTTCCACTCAACTTTAACCCg |
| 9 | gTCCCTgCCTCTATATCTTTCAGGGAACCATTTCCTGTTGTTCAG | 10 | GGTAGTCCCTCACATCCTCAGGTTCTTCCACTCAACTTTAACCCg |
| 11 | gTCCCTgCCTCTATATCTTTAGGCCTCTGGCTTGCAGGTACAGGA | 12 | AGGTGCTGTTGGATGGTCTTCACAGTTCCACTCAACTTTAACCCg |
| 13 | gTCCCTgCCTCTATATCTTTCCTGTGCAGCATGTTGAGCTGGCCC | 14 | GTCAGAAGGGCGAGGCAGGCCAGATTTCCACTCAACTTTAACCCg |
| 15 | gTCCCTgCCTCTATATCTTTTCATCACCAGGGACACAGCATTGGA | 16 | CATCCACATTCTCCTTTCTGATTCTTTCCACTCAACTTTAACCCg |
| 17 | gTCCCTgCCTCTATATCTTTAGGGCCTGCTTGGCTCTCTCCCCAG | 18 | TGGTCAAAGTCAGTGCGTTCAAAGGTTCCACTCAACTTTAACCCg |
| 19 | gTCCCTgCCTCTATATCTTTAGAGTTCTCCATCAGGGATCTGACT | 20 | CAGGTTCCTGATGTCCTGGCATCTGTTCCACTCAACTTTAACCCg |
| 21 | gTCCCTgCCTCTATATCTTTTGTTGTAGGCAATGCCCAGGAAGGC | 22 | TGGCAATTTCGGCAATGCGCAGCAGTTCCACTCAACTTTAACCCg |
| 23 | gTCCCTgCCTCTATATCTTTCGGGAGATGTCCTTCACTCTGATTC | 24 | TGGATCAGCATTCTCCCACCATCGGTTCCACTCAACTTTAACCCg |
| 25 | gTCCCTgCCTCTATATCTTTCACCAGGGTCTTGGTCCTGCCAATG | 26 | CAGGGCCTTCTCCACACCAGCTGTGTTCCACTCAACTTTAACCCg |
| 27 | gTCCCTgCCTCTATATCTTTCCACCAGCTTGGTAACCCCCAGGGA | 28 | CCACACCAGACACAGAGATCCATCTTTCCACTCAACTTTAACCCg |
| 29 | gTCCCTgCCTCTATATCTTTCATAGATCAGGCGGTGGGTGGCCTC | 30 | ATCTCTGCCCAGAGTCATCCTTGGCTTCCACTCAACTTTAACCCg |
| 31 | gTCCCTgCCTCTATATCTTTGCAGAGTGGCCAGACCAGGCCAGGT | 32 | GCCATGTCCCTGGCAGCACCCACTCTTCCACTCAACTTTAACCCg |
| 33 | gTCCCTgCCTCTATATCTTTTTCAGGGATGGACACACCAGCCCTG | 35 | ATTGGTCCAGCCACCAGCCTGCATGTTCCACTCAACTTTAACCCg |
| 35 | gTCCCTgCCTCTATATCTTTTGATGTAGTTCATCACTATGTTCAC | 36 | TGGCCCCAGTCTCAGAGTCCAGGTTTTCCACTCAACTTTAACCCg |

**Supplementary Table 2**. Statistical analysis of the data presented in Figures 2-7. Partial Eta^2^ = proportion of explained variance. Dorsal hippocampus, dHipp; Dorsal striatum, DS; Fixed ratio, FR; Fluorescence in situ hybridization chain reaction, HCR FISH; Nucleus accumbens, NAc; Self-administration, SA

Figure 2: males only

| **Figure number** | **Factor name** | **F value** | **P value** | **Partial Eta^2^** |
| --- | --- | --- | --- | --- |
| 2A: HCR FISH analysis: *Oprm1* mRNA NAc | Between-subjects: Genotype | F(1,8) = 3.0 | 0.123 | 0.270 |
| 2A: HCR FISH analysis: *Oprm1* mRNA DS | Between-subjects: Genotype | F(1,8) = 3.9 | 0.083 | 0.330 |
| 2A: HCR FISH analysis: *Oprm1* mRNA dHipp | Between-subjects: Genotype | F(1,8) = 0.5 | 0.480 | 0.064 |
| 2B: HCR FISH analysis: Cre +/- NAc | Within subjects: Cre +/- | F(1,4) = 356.1 | **<0.001*** | 0.989 |
| 2B: HCR FISH analysis: Cre +/- DS | Within subjects: Cre +/- | F(1,4) = 652.4 | **<0.001*** | 0.994 |
| 2B: HCR FISH analysis: Cre +/- dHipp | Within subjects: Cre +/- | F(1,4) = 172.3 | **<0.001*** | 0.977 |

Figure 3: males and females

| **Figure number** | **Factor name** | **F value** | **P value** | **Partial Eta^2^** |
| --- | --- | --- | --- | --- |
| 3A: NAc [^35^S]GTPγS activity | Between subjects: Genotype | F(1,10) = 2.1 | 0.178 | 0.173 |
| 3A: DS [^35^S]GTPγS activity | Between subjects: Genotype | F(1,10) = 0.2 | 0.636 | 0.023 |
| 3B: NAc [^3^H]DAMGO binding | Between subjects: Genotype | F(1,10) = 0.05 | 0.820 | 0.005 |
| 3B: DS [^3^H]DAMGO binding | Between subjects: Genotype | F(1,10) = 6.7 | **0.027*** | 0.400 |
| 3C: NAc [^3^H]DAMGO binding: NAc shell | Between-subjects: Genotype  Within-subjects: Lesion (Vehicle, Caspase3)  Genotype x Lesion | F(1,9) = 3.6  F(1,9) = 12.5  F(1,9) = 9.9 | 0.089  **0.006***  **0.012*** | 0.288  0.581  0.525 |
| 3C: NAc [^35^S]GTPγS activity: NAc shell | Between-subjects: Genotype  Within-subjects: Lesion (Vehicle, Caspase3)  Genotype x Lesion | F(1,9) = 1.7  F(1,9) = 5.9  F(1,9) = 1.6 | 0.230  **0.038***  0.236 | 0.156  0.396  0.152 |
| 3D: RNAscope: NAc shell | Between-subjects: Genotype  Within-subjects: Lesion (Vehicle, Caspase3)  Genotype x Lesion | F(1,9) = 0.2  F(1,9) = 11.3  F(1,9) = 8.4 | 0.666  **0.008***  **0.018*** | 0.022  0.556  0.482 |
| 3D: RNAscope: NAc core | Between-subjects: Genotype  Within-subjects: Lesion (Vehicle, Caspase3)  Genotype x Lesion | F(1,9) = 1.7  F(1,9) = 3.2  F(1,9) = 4.8 | 0.224  0.106  0.056 | 0.159  0.264  0.348 |

Figure 4: males and females

| **Figure number** | **Factor name** | **F value** | **P value** | **Partial Eta^2^** |
| --- | --- | --- | --- | --- |
| 4A: Acquisition: Pellets | Between-subjects: Genotype  Within-subjects: Session  Genotype x Session | F(1,18) = 0.04  F(6,108) = 24.1  F(6,108) = 0.5 | 0.839  **<0.001***  0.833 | 0.002  0.573  0.025 |
| 4A: Acquisition: Lever presses | Between-subjects: Genotype  Within-subjects: Session  Genotype x Session | F(1,18) = 0.1  F(6,108) = 7.8  F(6,108) = 0.3 | 0.711  **<0.001***  0.923 | 0.008  0.303  0.018 |
| 4A: FR response: Pellets | Between-subjects: Genotype  Within-subjects: FR requirement  Genotype x FR requirement | F(1,18) = 0.0  F(4,72) = 34.4  F(4,72) = 1.4 | 0.987  **<0.001***  0.248 | 0.000  0.656  0.071 |
| 4A: FR response: Lever presses | Between-subjects: Genotype  Within-subjects: FR requirement  Genotype x FR requirement | F(1,18) = 0.03  F(4,72) = 12.3  F(4,72) = 1.2 | 0.874  **<0.001***  0.311 | 0.001  0.407  0.063 |
| 4B: Heroin SA: infusions | Between-subjects: Genotype  Within-subjects 1: Heroin Dose  Within-subjects 2: Session  Genotype x Session  Heroin Dose x Genotype  Heroin Dose x Session  Genotype x Heroin Dose x Session | F(1,25) = 0.1  F(1,25) = 62.2  F(5,125) = 5.5  F(5,125) = 0.9  F(1,25) = 0.9  F(5,125) = 10.0  F(5,125) = 0.5 | 0.767  **<0.001***  **<0.001***  0.515  0.365  **<0.001***  0.768 | 0.004  0.713  0.181  0.033  0.033  0.286  0.020 |
| 4B: Heroin: Extinction responding | Between-subjects: Genotype  Within-subjects: Session (1-7)  Genotype x Session | F(1,25) = 1.3  F(6,150) = 49.9  F(6,150) = 0.1 | 0.273  **<0.001***  0.994 | 0.048  0.666  0.005 |
| 4B: Heroin: Context-induced reinstatement | Between-subjects: Genotype  Within-subjects: Context (A, B)  Genotype x Context | F(1,25) = 0.00  F(1,25) = 65.4  F(1,25) = 0.1 | 0.963  **<0.001***  0.742 | 0.000  0.723  0.004 |
| 4B: Heroin: Reacquisition | Between-subjects: Genotype  Within-subjects: Session Hour (1-6)  Genotype x Session Hour | F(1,25) = 0.5  F(5,125) = 6.0  F(5,125) = 1.6 | 0.474  **<0.001***  0.166 | 0.021  0.194  0.060 |

Figure 5: males and females

| **Figure number** | **Factor name** | **F value** | **P value** | **Partial Eta^2^** |
| --- | --- | --- | --- | --- |
| 5A. von Frey: dose response | Between subjects: Genotype  Within subjects: Dose  Genotype x Dose | F(1,22.35) = 0.0  F(3,47.72) = 73.0  F(3,47.72) = 0.1 | 0.931  **<0.001***  0.931 |  |
| 5A. von Frey: morphine + naloxone | Between subjects: Genotype  Within subjects: Condition  Genotype x Condition | F(1,17.45) = 0.0  F(2,17.45) = 65.4  F(2,17.45) = 0.1 | 0.917  **<0.001***  0.924 |  |
| 5A. von Frey: morphine + tolerance | Between subjects: Genotype  Within subjects: Condition  Genotype x Condition | F(1,30.54) = 0.0  F(2,30.54) = 87.4  F(2,30.54) = 0.1 | 0.897  **<0.001***  0.905 |  |
| 5B. Tail flick:  dose response | Between subjects: Genotype  Within subjects: Dose  Genotype x Dose | F(1,19.4) = 0.0  F(4,61.58) = 58.1  F(4,61.58) = 0.6 | 0.979  **<0.001***  0.700 |  |
| 5B. Tail flick:  morphine + naloxone | Between subjects: Genotype  Within subjects: Condition  Genotype x Condition | F(1,19.75) = 0.1  F(2,23.01) = 22.8  F(2,23.01) = 0.1 | 0.723  **<0.001***  0.908 |  |
| 5B. Tail flick:  morphine + tolerance | Between subjects: Genotype  Within subjects: Condition  Genotype x Condition | F(1,18.93) = 0.2  F(2,33.63) = 20.8  F(2,33.63) = 0.1 | 0.658  **<0.001***  0.885 |  |
| 5C. Lactic acid dose response: change baseline | Between subjects: Genotype  Within subjects: Dose  Genotype x Dose | F(1,13) = 0.02  F(3,39) = 17.2  F(3,39) = 0.9 | 0.889  **<0.001***  0.471 | 0.002  0.569  0.062 |
| 5C. Morphine dose response: change baseline | Between subjects: Genotype  Within subjects: Dose  Genotype x Dose | F(1,13) = 0.4  F(3,39) = 58.1  F(3,39) = 0.05 | 0.557  **<0.001***  0.984 | 0.027  0.817  0.004 |
| 5C. Morphine + lactic acid: change baseline | Between subjects: Genotype  Within subjects: Dose  Genotype x Dose | F(1,13) = 1.7  F(2,26) = 25.0  F(2,26) = 0.2 | 0.217  **<0.001***  0.828 | 0.115  0.658  0.014 |

Figure 6: males and females

| **Figure number** | **Factor name** | **F value** | **P value** | **Partial Eta^2^** |
| --- | --- | --- | --- | --- |
| 6A: Food Acquisition: Pellets | Between-subjects 1: Genotype  Between-subjects 2: Sex  Genotype x Sex  Within-subjects: Session  Genotype x Session  Sex x Session  Genotype x Sex x Session | F(1,51) = 0.1  F(1,51) = 1.2  F(1,51) = 0.3  F(11,561) = 24.2  F(11,561) = 1.7  F(11,561) = 1.6  F(11,561) = 0.7 | 0.753  0.27  0.605  **<0.001***  0.072  0.109  0.727 | 0.002  0.024  0.005  0.322  0.032  0.030  0.014 |
| 6A: Food FR requirement: pellets | Between-subjects 1: Genotype  Between-subjects 2: Sex  Genotype x Sex  Within-subjects: FR requirement  Genotype x FR requirement  Sex x FR requirement  Genotype x Sex x FR requirement | F(1,22) = 1.0  F(1,22) = .001  F(1,22) = 0.2  F(6,132) = 49.8  F(6,132) = 0.6  F(6,132) = 1.7  F(6,132) = 0.5 | 0.339  0.981  0.665  **<0.001***  0.761  0.133  0.773 | 0.042  0.000  0.009  0.693  0.025  0.071  0.024 |
| 6B. Heroin Acquisition: dose response | Between-subjects 1: Genotype  Between-subjects 2: Sex  Genotype x Sex  Within-subjects 1: Dose  Dose x Sex  Dose x Genotype  Dose x Sex x Genotype | F(1,49) = 5.1  F(1,49) = 4.1  F(1,49) = 1.9  F(3,147) = 0.7  F(3,147) = 1.5  F(3,147) = 0.9  F(3,147) = 2.1 | **0.029***  **0.049***  0.169  0.545  0.225  0.437  0.109 |  |
| 6C. Heroin Acquisition 3 day mean: dose response | Between-subjects 1: Genotype  Between-subjects 2: Sex  Genotype x Sex  Within-subjects: Dose  Sex x Dose  Genotype x Dose  Sex x Genotype x Dose | F(1,49) = 5.1  F(1,49) = 4.1  F(1,49) = 2.0  F(3,147) = 0.7  F(3,147) = 1.5  F(3,147) = 0.9  F(3,147) = 2.0 | **0.029***  **0.049***  0.167  0.547  0.224  0.437  0.111 | 0.094  0.077  0.039  0.014  0.029  0.018  0.040 |
| 6D. Heroin Maintenance: within-session dose response | Between-subjects 1: Genotype  Between subjects 2: Sex  Genotype x Sex  Within-subjects: Dose  Dose x Genotype  Dose x Sex  Dose x Genotype x Dose | F(1,46) = 4.7  F(1,46) = 3.0  F(1,46) = 0.1  F(3,138) = 88.2  F(3,138) = 3.1  F(3,138) = 0.8  F(3,138) = 1.2 | **0.035***  0.088  0.714  **<0.001***  **0.029***  0.502  0.301 | 0.093  0.062  0.003  0.657  0.063  0.017  0.026 |
| 6E. Heroin Maintenance: within-session FR response | Between-subjects 1: Genotype  Between subjects 2: Sex  Genotype x Sex  Within-subjects: FR requirement  FR requirement x Genotype  FR requirement x Sex  FR requirement x Genotype x Sex | F(1,46) = 14.0  F(1,46) = 3.0  F(1,46) = 0.8  F(6,276) = 54.6  F(6,276) = 1.7  F(6,276) = 0.6  F(6,276) = 1.0 | **<0.001**  0.088  0.387  **<0.001***  0.130  0.741  0.395 | 0.234  0.062  0.016  0.543  0.035  0.013  0.022 |
| 6F. Heroin Maintenance: extended access | Between-subjects 1: Genotype  Between subjects 2: Sex  Genotype x Sex  Within-subjects: Hour  Hour x Genotype  Hour x Sex  Hour x Genotype x Sex | F(1,46) = 5.4  F(1,46) = 8.0  F(1,46) = 0.03  F(8,365) = 5.1  F(8,365) = 2.0  F(8,365) = 1.7  F(8,365) = 1.6 | **0.025***  **0.007***  0.870  **<0.001***  **0.041***  0.100  0.112 |  |
| 6G: NAc [^3^H]DAMGO binding | Between-subjects: Genotype | F(1,24) = 11.0 | **0.003*** | 0.315 |

Figure 6: males only

| **Figure number** | **Factor name** | **F value** | **P value** | **Partial Eta^2^** |
| --- | --- | --- | --- | --- |
| 6A: Food Acquisition: Pellets | Between-subjects 1: Genotype  Within-subjects: Session  Session x Genotype | F(1,26) = 0.4  F(11,286) = 14.3  F(11,286) = 1.4 | 0.528  **<0.001***  0.182 | 0.016  0.355  0.050 |
| 6A: Food FR requirement: pellets | Between-subjects: Genotype  Within-subjects: FR requirement  Genotype x FR requirement | F(1,13) = 1.2  F(6,78) = 21.8  F(6,78) = 0.6 | 0.298  **<0.001***  0.712 | 0.894  0.627  0.046 |
| 6B. Heroin Acquisition: dose response | Between-subjects: Genotype  Within-subjects 1: Dose  Dose x Genotype | F(1,24) = 8.0  F(3,72) = 1.0  F(3,72) = 1.5 | **0.009***  0.418  0.209 |  |
| 6C. Heroin Acquisition 3 day mean: dose response | Between-subjects: Genotype  Within-subjects: Dose  Dose x Genotype | F(1,24) = 8.0  F(3,72) = 1.0  F(3,72) = 1.5 | **0.009***  0.420  0.214 | 0.250  0.038  0.060 |
| 6D. Heroin Maintenance: within-session dose response | Between-subjects: Genotype  Within-subjects: Dose  Dose x Genotype | F(1,22) = 4.2  F(3,66) = 49.4  F(3.66) = 4.6 | 0.053  **<0.001***  **0.006*** | 0.160  0.692  0.172 |
| 6E. Heroin Maintenance: within-session FR response | Between-subjects: Genotype  Within-subjects: FR requirement  FR requirement x Genotype | F(1,22) = 5.2  F(6,132) = 40.6  F(6,132) = 0.4 | **0.033***  **<0.001***  0.854 | 0.191  0.649  0.019 |
| 6F. Heroin Maintenance: extended access | Between-subjects: Genotype  Within-subjects: Hour  Hour x Genotype | F(1,22.03) = 2.6  F(8,174.1) = 2.0  F(8,174.1) = 0.6 | 0.122  0.053  0.739 |  |

Figure 6: females only

| **Figure number** | **Factor name** | **F value** | **P value** | **Partial Eta^2^** |
| --- | --- | --- | --- | --- |
| 6A: Food Acquisition: Pellets | Between-subjects 1: Genotype  Within-subjects: Session  Session x Genotype | F(1,25) = 0.02  F(11,275) = 11.3  F(11,275) = 1.0 | 0.894  **<0.001***  0.443 | 0.001  0.312  0.039 |
| 6A: Food FR requirement: pellets | Between-subjects: Genotype  Within-subjects: FR requirement  Genotype x FR requirement | F(1,9) = 0.1  F(6,54) = 30.9  F(6,54) = 0.5 | 0.729  **<0.001***  0.785 | 0.014  0.775  0.055 |
| 6B. Heroin Acquisition: dose response | Between-subjects: Genotype  Within-subjects 1: Dose  Dose x Genotype | F(1,25) = 0.3  F(3,75) = 1.2  F(3,75) = 1.5 | 0.578  0.324  0.228 |  |
| 6C. Heroin Acquisition 3 day mean: dose response | Between-subjects: Genotype  Within-subjects: Dose  Dose x Genotype | F(1,25) = 0.3  F(3,75) = 1.2  F(3,75) = 1.5 | 0.579  0.323  0.230 | 0.013  0.045  0.056 |
| 6D. Heroin Maintenance: within-session dose response | Between-subjects: Genotype  Within-subjects: Dose  Dose x Genotype | F(1,25) = 2.0  F(3,75) = 40.0  F(3,75) = 0.4 | 0.168  **<0.001***  0.719 | 0.075  0.616  0.018 |
| 6E. Heroin Maintenance: within-session FR response | Between-subjects: Genotype  Within-subjects: FR requirement  FR requirement x Genotype | F(1,24) = 9.1  F(6,144) = 20.5  F(6,144) = 2.0 | **0.006***  **<0.001***  0.073 | 0.276  0.461  0.076 |
| 6F. Heroin Maintenance: extended access | Between-subjects: Genotype  Within-subjects: Hour  Hour x Genotype | F(1,24.02) = 2.8  F(8,191) = 4.4  F(8,191) = 2.6 | 0.105  **<0.001***  **0.010*** |  |

Figure 7: males and females

| **Figure number** | **Factor name** | **F value** | **P value** | **Partial Eta^2^** |
| --- | --- | --- | --- | --- |
| 7A: Food Acquisition: Pellets | Between-subjects 1: Genotype  Within-subjects: Session  Session x Genotype | F(1,26) = 0.01  F(11,286) = 6.8  F(11,286) = 0.7 | 0.913  **<0.001***  0.729 | 0.000  0.208  0.027 |
| 7A: Food FR requirement: pellets | Between-subjects: Genotype  Within-subjects: FR requirement  Genotype x FR requirement | F(1,26) = 0.7  F(6,156) = 50.7  F(6,156) = 0.2 | 0.420  **<0.001***  0.980 | 0.025  0.661  0.007 |
| 7B. Heroin Acquisition: dose response | Between-subjects: Genotype  Within-subjects 1: Dose  Within-subjects 2: Session  Dose x Genotype  Dose x Session  Session x Genotype  Dose x Genotype x Session | F(1,26) = 2.4  F(3,78) = 4.7  F(2,52) = 4.6  F(3,78) = 1.7  F(6,156) = 2.5  F(2,52) = 1.1  F(6,156) = 1.1 | 0.132  **0.005***  **0.014***  0.168  **0.023***  0.335  0.380 | 0.085  0.153  0.152  0.062  0.089  0.041  0.040 |
| 7C. Heroin Acquisition: 3-day mean dose response | Between-subjects: Genotype  Within-subjects 1: Dose  Dose x Genotype | F(1,26) = 0.001  F(3,78) = 7.0  F(3,78) = 0.7 | 0.972  **<0.001***  0.576 | 0.000  0.212  0.025 |
| 7D. Heroin Maintenance: within-session dose response | Between-subjects: Genotype  Within-subjects: Dose  Dose x Genotype | F(1,24) = 0.6  F(3,72) = 49.1  F(3,72) = 0.3 | 0.449  **<0.001***  0.816 | 0.024  0.672  0.013 |
| 7E. Heroin Maintenance: within-session FR response | Between-subjects 1: Genotype  Within-subjects: FR requirement  FR requirement x Genotype | F(1,24) = 1.0  F(6,144) = 16.4  F(6,144) = 0.3 | 0.325  **<0.001***  0.922 | 0.040  0.405  0.013 |
| 7F. Heroin Maintenance: extended access | Between-subjects 1: Genotype  Within-subjects: Hour  Hour x Genotype | F(1,24) = 1.9  F(8,192) = 10.6  F(8,192) = 0.4 | 0.182  **<0.001***  0.939 | 0.073  0.306  0.015 |
